# Recurrent extreme climatic events are driving gorgonian populations to local extinction: low adaptive potential to marine heatwaves

**DOI:** 10.1101/2024.05.13.593802

**Authors:** Sandra Ramirez-Calero, D Gómez-Gras, A Barreiro, N Bensoussan, L Figuerola-Ferrando, M Jou, A López-Sanz, P López-Sendino, A Medrano, I Montero-Serra, M Pagès-Escolà, C Linares, JB Ledoux, J Garrabou

## Abstract

Extreme climatic events (ECEs), such as marine heatwaves (MHWs), are a major threat to biodiversity. Understanding the variability in ecological responses to recurrent ECEs within species and underlying drivers arise as a key issue owing to their implications for conservation and restoration. Yet, our knowledge on such ecological responses is limited since it has been mostly gathered following “single-event approaches” focused on one particular event. These approaches provide snapshots of ecological responses but fall short of capturing heterogeneity patterns that may occur among recurrent ECEs, questioning current predictions regarding biodiversity trends. Here, we adopt a “multi-event” perspective to characterize the effects of recurrent ECEs and the ecological responses in *Paramuricea clavata*, a Mediterranean temperate coral threatened by MHWs. Through a common-garden experiment repeated three consecutive years with the same individuals from three populations, we assessed the respective roles of environmental (year effect), genetic (population effect) and phenotypic (*population-by-environment* interactions effect) components in the ecological response to recurrent heat stress. The environmental component (year) was the main driver underlying the responses of *P. clavata* colonies across experiments. To build on this result, we showed that: i) the ecological responses were not related to population (genetic isolation) and individual (multilocus heterozygosity) genetic make-up, ii) while all the individuals were characterized by a high environmental sensitivity (*genotype-by-environment* interactions) likely driven by *in-situ* summer thermal regime. We confront our experimental results to *in situ* monitoring of the same individuals conducted in 2022 following two MHWs (2018, 2022). This confirms that the targeted populations harbor limited adaptive and plastic capacities to on-going recurrent ECEs and that *P. clavata* might face unavoidable population collapses in shallow Mediterranean waters. Overall, we suggest that biodiversity forecasts based on “single event” experiments may be overly optimistic and underscore the need to consider the recurrence of ECEs in assessing threats to biodiversity.

## Introduction

Extreme climate events (ECE) linked to climate change such as heatwaves pose significant challenges for biodiversity (Jacox et al., 2020; Maxwell et al., 2019; Parmesan et al., 2000; Pinsky et al., 2019; Ummenhofer & Meehl, 2017). ECEs have been associated with the increased frequency of mass mortality events (MMEs), accelerating species demographic decline (Leung *et al*., 2017; Smale et al., 2013) and questioning the future of biodiversity (Smale et al. 2019). Besides demographic decline, field surveys revealed heterogeneity in the patterns of ecological responses to ECEs across taxonomic (species, populations, and individuals), spatial (regions) and temporal (years) scales (van Bergen et al., 2020; Hughes et al., 2003; Pansch et al., 2018). Yet, to date, the potential effects of recurrent ECEs and the related temporal variability in ecological responses has been poorly considered (but see Ahrens et al., 2021; Brown et al., 2023; Brown & Barott, 2022; Husson et al., 2022; Logan et al., 2014; Regan & Sheldon, 2023). Indeed, most of our knowledge on ecological responses to ECEs has been gathered studying one particular event following a “single-event approach” (Altwegg et al., 2017). This approach provides a snapshot of the ecological responses but falls short of revealing the consequences of recurrent ECEs on biodiversity. Consequently, the need to develop a “multi-events” perspective has been recently acknowledged to improve predictions on species abilities to face ECEs (Bailey & Van De Pol, 2016).

Ecological responses to ECEs, considered here as phenotypes, are shaped by the interplay among ‘genetic’, ‘environmental’ and ‘plastic’ components (Merilä & Hendry, 2014). This interplay has been investigated using common garden experiments (Malyshev et al., 2016) and long-term time series (Regan & Sheldon, 2023). The ‘genetic’ component relies on the standing genetic variation (Dixon et al., 2015), shaped by the balance between evolutionary forces (*e.g.* local adaptation, genetic drift; Bay & Palumbi, 2014). The ‘environmental’ component includes biotic (*e.g.* species interactions) and abiotic (*e.g.* temperature) factors that characterize a specific habitat and influence ecological responses (Scheiner, 1993). Finally, ‘plastic’ components result from the interaction of genetic and environmental components. This *“population-by-environment”* and “*genotype-by-environment*” interactions underlie plasticity in phenotypic response at the population and individual levels (Matesanz et al., 2019). Individual phenotypic plasticity, considered as a single genotype expressing different phenotypes function of the environment, can provide a short-term buffer allowing organisms to immediately face ECEs before genetic adaptation occurs (Chevin et al., 2010; Reusch, 2014).

Marine heatwaves (MHWs), known as discrete periods of anomalously warm water (Frölicher et al., 2018; Smith et al., 2023), are some of the most challenging ECEs for marine diversity, threatening ecosystem’s structure and functioning (Smale et al., 2019) from kelp forests (Arafeh-Dalmau et al., 2020) to coral reefs (Genin et al., 2020; Gómez-Gras et al., 2021a; Hughes et al., 2021). In the last two decades, the Mediterranean Sea has been recurrently impacted by MHWs driving mass mortality events (MMEs) impacting multiple phyla of benthic macro-invertebrates (Cramer et al., 2018; Garrabou et al., 2022). In this particular case, as in other marine ecosystems, field surveys (Garrabou et al., 2001) and long-term monitoring studies (Gómez-Gras et al., 2021b; Montero-Serra et al., 2019) combined with “single event” experiments in controlled conditions (Crisci et al., 2017; Gomez-Gras et al. 2022), provided insights into the heterogeneity of ecological responses within species impacted by MHWs. Yet, whatever the system considered, the impact of genetic, environmental, and plastic components underlying the differential ecological responses remain poorly understood, particularly, in the context of recurrent MHWs (but see Hughes et al., 2021).

We aim to advance the characterization of the impacts of recurrent ECEs on within-species diversity and to infer the respective roles of the genetic, environmental, and plastic components in the ecological responses. We adopt a multi-events perspective combining experiments and *in-situ* survey focusing on the Mediterranean scene and on the red gorgonian *Paramuricea clavata* (Risso, 1826). This habitat-forming octocoral is a well-suitable candidate given the impacts of MHWs on shallow populations monitored for 20 years (Garrabou et al., 2021, see below). We repeated during three consecutive years (2015, 2016, 2017) a common-garden thermal stress experiment at fine spatial scale (populations separated by < 1 km), in which we controlled for different aspects of genetic (same genotypes tested) and environmental (same experimental set-up) components. Specifically, we; i) estimated the percentage of heterogeneity in the ecological responses, respectively explained by the taxonomic (individual and populations) and temporal (years) variabilities; ii) tested the significance of the genetic (population effect), environmental (year effect) and plastic (population-by-years effect) components on the ecological responses. We discussed the obtained results in the light of iii) the populations and individuals’ genetic make-up (*i.e.* genetic drift and heterosis); iv) environmental sensitivity analyses looking at genotype-by-environment interactions and v) *in-situ* summer thermal regimes. We vi) contrasted the ecological responses from the experiments to the ecological responses reported from a field survey of the same individuals conducted in 2022 after two MHWs.

Compared to previously published single event approaches, our results point toward a lack of adaptive potential, whether plastic or genetic, of the shallow populations of *P. clavata* to the recurrent MHWs. We suggest to cautiously consider predictions of biodiversity trends based on “single event” approaches.

## Materials and methods

### Species of interest

The red gorgonian *P. clavata* is a habitat-forming octocoral with an important role in the structure and functioning of the Mediterranean coralligenous habitats (Boavida et al., 2016; Gómez-Gras et al., 2021a; Ponti et al., 2014, 2018). This species displays low population dynamics with slow growth, recruitment, and recovery rates (Gómez-Gras et al., 2021a; Gómez-Gras et al., 2021b), as well as restricted dispersal abilities with a highly reduced resilience (Coma et al., 1998; Ledoux et al., 2018; Linares et al., 2008; Mokhtar-Jamaï et al., 2011). *Paramuricea clavata* has been particularly impacted by recurrent marine heat waves in the past 20 years (Cebrian et al., 2012; Garrabou et al., 2021; Gómez-Gras et al., 2021b). It was included in the IUCN red list of vulnerable Mediterranean Anthozoans (Otero et al., 2017). “Single event” experiments using common garden set-ups identified the thermal risk zone of this species for temperatures over 25° C (Crisci et al 2017). These experiments have characterized ecological responses and underlying processes in populations from contrasted environments (depth) and separated by distances of tens to thousands of kms (Arizmendi-Mejía et al., 2015b; Crisci et al., 2017). Altogether, the ecological importance of *P. clavata* as a habitat-forming species and the extensive available ecological information make this model highly relevant to evaluate the roles of the genetic, environmental, and plastic components on the ecological responses to recurrent ECEs.

### Identification of the colonies

Ninety adult colonies (>50 cm) were randomly chosen around 15 m depth from three different sites separated by hundreds of meters at Medes Islands, Spain (42°02’60.00” N 3°12’60.00” E): Pota del Llop (N=30), La Vaca (N=30) and Tascons (N=30, Figure 1a). Each colony was marked during September of 2014 using plastic tags with a unique ID (Figure 1b). From every marked colony, two apical fragments of 10 cm were collected between the 21^st^ of September and 22^nd^ of October of 2015, 2016 and 2017. These fragments were retrieved in 2 L bags of water and immediately transported (maximum transportation time 2h) in coolers to the Aquarium Experimental Zone (ZAE) of the Institut de Ciències del Mar (ICM-CSIC, Barcelona, Spain).

**Figure 1.**
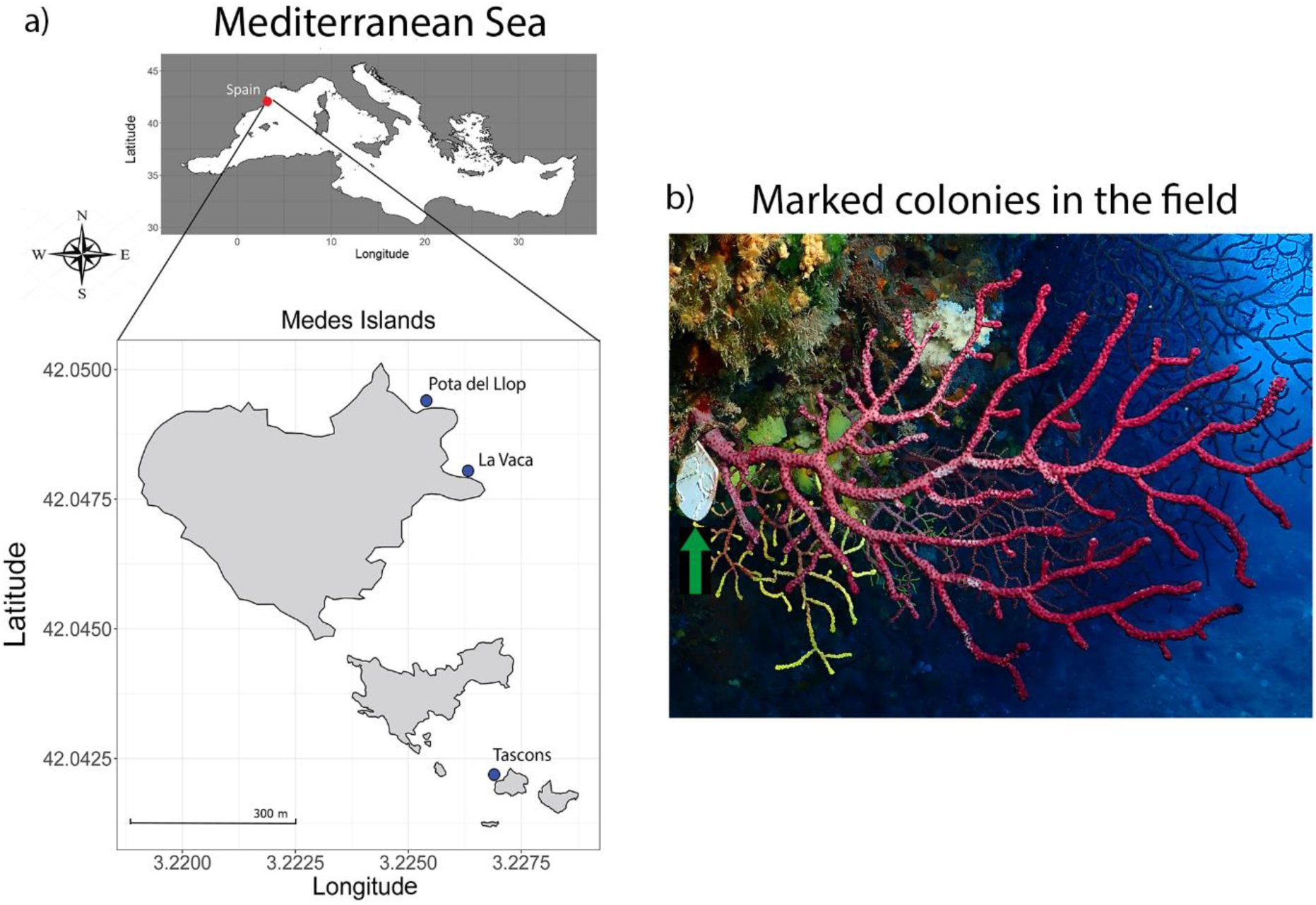
a) *Paramuricea clavata* sampling sites at Medes Islands, Spain, b) Tagged colony of *P. clavata* in Tascon’s location (green arrow).

### Common garden set-up repeated in 2015, 2016 and 2017

Upon arrival, colony fragments were mechanically fixed to experimental plates as described in Crisci et al. (2017) (Figure S1). All colonies were acclimated for one week in an open water system with 50 *μ*m sand-filtered running seawater at 17-18°C. No mortality signs and tissue necrosis were detected during this period in any of the colonies and/or years. Moreover, all sampled colonies showed active polyps during feeding. After the acclimation period, we conducted the common garden experiments as described in Crisci et al. (2017). Specifically, each fragment was divided into two fragments, one for control (18°C) and one for the heat stress treatment (25°C). For the heat stress treatment, temperature was increased from 18 to 25 °C over a period of 3 days and maintained at 25°C for the next 28 days. Colonies were fed during the entire experimental course (see Supplementary information for further details about the set up).

### Phenotypic response: survival analysis, individual fitness, and modeling the response to thermal stress

Several descriptors were then used to statistically compare the ecological responses (differences in thermotolerance) among colonies and populations. The percentage of tissue necrosis (extent of injury) by colony was monitored visually every day. Ecological impacts on *P. clavata* were considered as “low” when 10-30% necrosis was observed, “moderate” when necrosis was >30-60% and “severe” for >60% necrosis, following the impact classification of the T-MEDNet mass mortality database (e.g. Garrabou et al., 2019). In addition, we estimated for each population and year: i) the daily extent of injury per colony (% of tissue necrosis), ii) the daily percentage of affected colonies (>10% of tissue necrosis), and iii) the daily percentage of dead colonies (100% tissue necrosis).

To model the response to thermal stress across years and populations, we computed a Principal Component Analysis (PCA) with individuals and years as cases and the percentage of necrosis per day as variables. A first exploratory PCA explained 50.29% of the variance and was strongly correlated with the percentage of necrosis from day 10^th^ to 28^th^ (Figure S2a). To improve the fitness of the data, days 1 to 9 were removed from the data set. Thus, the second PCA explained 75% of the variance for the remaining days (Figure S2b). All PCAs were performed with *PCA()* function from *FactoMineR* R package (R Core Team, 2022; Lê et al., 2008). Scores from the second PCA were considered as a proxy for individual fitness as they were correlated to the baseline percentages of necrosis. These scores were employed as response variable for two linear models: 1) including ‘population’ and ‘year’ as predictor variables (*i.e.* fixed factors); 2) using ‘individual’ as a random factor, added to the previous predictor variables, and affecting only the model intercepts. Available data were not sufficient to fit all the extra number of parameters (*i.e.* coefficients), thus the effect of the factor ‘individual’ was not tested in the slope of the model. We consider the ‘individual’ effect in the intercept as a different baseline of resistance to thermal stress. Linear models were fitted using *lm()* and *lmer()* functions from the *stats* and the *lme4()* R*-*packages (Fox & Weisberg, 2019). To meet the assumptions of residual normality and homoscedasticity, we transformed the response variable with the Box-Cox transformation, implemented with the *boxcox()* function from *MASS* R package (Venables & Ripley, 2002). These assumptions were tested with the Shapiro-Wilks and Levene tests using *Shapiro.test()* and *leveneTest()* from *car* R-package (Fox & Weisberg, 2019). The best model was selected based on AIC (Akaike information criterion) using the function *anova()* (R Core Team, 2022) and a *post-hoc* Tukey test with the *glht()* function from *multcomp* R-package (Hothorn et al., 2008). We estimated the percentage of the contribution of each factor to the total variance computing the difference in log-likelihood between models with and without each factor (individual, population, and year) pondered by the degrees of freedom. The random intercepts obtained for every individual in the linear model 2 (hereafter *“nec-int*”) were used as a proxy for individual fitness to test for heterosis (see below).

### Genetic components: population structure and individual heterosis

DNA extractions, genotyping protocols, microsatellite characteristics, quality check, and analyses of genetic diversity are described in Supplementary information. We characterized the genetic diversity and structure of the three sampling sites by genotyping 87 individuals collected in at least two of the three years with 14 microsatellites.

We conducted a Discriminant Analysis of Principal Components (DAPC; Jombart et al., 2010) from *adegenet* R-package (Jombart et al., 2010), by implementing the function *find_clusters()* and specifying a maximum number of clusters with *max.n.clust* = 3. A maximum number of 100 PCs was chosen and the lowest value of Bayesian Information Criterion (BIC) was applied to estimate the number of clusters. GENEPOP 4.1.4 (Rousset, 2008) was used to compute the overall and pairwise *F_STs_* estimators from Weir & Cockerham (1984). Genotypic differentiation between sites was tested using an exact test (Raymond & Rousset, 1995) with default parameters. In small and isolated populations, inbreeding depression can negatively impact individual fitness constraining the response to ECEs (Fitzpatrick & Reid, 2019). Accordingly, we estimated the genetic differentiation proper to each site by calculating the site-specific *F_ST_*and 95% High Probability Density Intervals in GESTE (Foll & Gaggiotti, 2006). The site-specific *F_ST_* estimates the relative impact of genetic drift on the differentiation of the considered site relative to the remaining ones.

At the individual level, a positive correlation between heterozygosity and fitness-related traits, known as heterosis, has been reported in some species (David, 1997). We looked for heterosis in the response to thermal stress testing the correlation between the values of *nec-int* as proxy for individual fitness and the standardized individual multilocus heterozygosities (*sMLH)* computed using the R package *InbreedR()* (Stoffel et al., 2016). The slope of the linear model [*lm(nec-int ~ sMLH)*] was compared to its null distribution obtained with a Monte-Carlo permutation test with 10,000 permutations.

### Environmental component: summer thermal environments

We analyzed and compared the *in situ* thermal regime at 15m depth experienced by the colonies before sampling. Temperature data for Medes islands was obtained from the T-MEDNet database, which follows a continuous temperature series since 2004 (von Schuckmann et al., 2019). We assessed the recent summer local thermal regime calculated over the 3-months period of June, July and August, prior to collection in September. We consider the timing and magnitude of the daily temperature cycles and the exposure to warm conditions; thus we considered the averages of maximum temperatures during the warmest periods of the year, number of days with high temperatures and the ecological threshold T23 (*i.e.* number of extreme heat days at ≥23°C). In addition, we considered positive temperature anomalies as the number of marine heat spikes (MHS) above the Inter-annual percentile 90^th^ (iT90 threshold) lasting less than five days, while prolonged discrete periods of anomalously warm water surpassing the iT90 percentile for more than 5 days, were considered as MHWs. Presence of MHWs were detected with Python module *marineHeatWaves* (Hobday et al., 2016), while impact categories registered at Medes islands were assigned following the classification of the T-MEDNet database (Garrabou et al., 2019; Hobday et al. 2018). MHWs categories were set as significant temperature foldchanges from the Inter-annual mean temperature and the iT90 threshold. Thus, “Low” classifications correspond to the temperature ranges between the inter-annual mean temperatures and iT90, while “Moderate” are considered as a 1-2-fold, “Strong” as a 2-3-fold and “Severe” as a 3-4-fold, respectively.

### Plastic component: ‘genotype-by-environment’ interactions and environmental sensitivity

We estimated the variability in the ‘*genotype-by-environment*’ interactions by characterizing the environmental sensitivity of each genotype following Falconer & Mackay (1996). We computed three environmental values corresponding to the yearly mean phenotypes (*i.e.* mean PCA scores over individuals for each year), considering the 66 genotypes present during the three years. We then plotted each individual phenotype (PCA score) against the environmental value for each year and we computed the regression slope, which is considered as an estimator of the environmental sensitivity of the genotype (Falconer & Mackay, 1996). We used this plot to classify the sensitivity of the genotypes in three categories adapted from Bonacolta et al. (2024). Resistant genotypes were expected to show low intercepts in the first year and approximated null slopes (*i.e.* low and constant level of necrosis in the 3 experiments). Hyper-sensitive genotypes were expected to show high intercepts in the first year and null slopes (*i.e.* high and constant level of necrosis in the 3 experiments). Finally, sensitive genotypes were expected to show low intercepts in the first year and positive slopes (*i.e.* increasing level of necrosis through time).

### Field survey of necrosis rates following 2018-2022 MMEs

Medes Islands were impacted by two MHWs in 2018 and 2022 with associated mass mortality events (Garrabou et al., 2022; Zentner et al., 2023). We surveyed by scuba diving the percentage of tissue necrosis *in situ* for the same individuals used in the experiments. This survey was done in October 2022. Differences in mean tissue necrosis of colonies between populations were tested using a parametric one-way ANOVA, followed by *post hoc* Tukey’s HSD tests.

## Results

### Phenotypic responses of P. clavata during the three common garden experiments

Signs of tissue necrosis were observed for all populations in the three years. In 2015 and 2016, the mean extent of injury was of moderate impact with values below the 60% at the end of the experiment (day 28^th^; mean ± SE): 32.14% ± 7.25 and 38.33% ± 6.67 for La Vaca; 27.03% ± 6.20 and 35.16% ± 7.09 for Pota del Llop and 37.89% ± 7.26 and 41.2% ± 6.91 for Tascons (Figure 2). On the contrary, in 2017, colonies showed severe impacts with >60% of average tissue necrosis earlier by day 14^th^ in all populations, and all colonies died by day 18^th^ (100% of tissue necrosis) with the exception of Pota del Llop which reached 100% of tissue necrosis by day 24^th^ (Figure 2).

**Figure 2.**
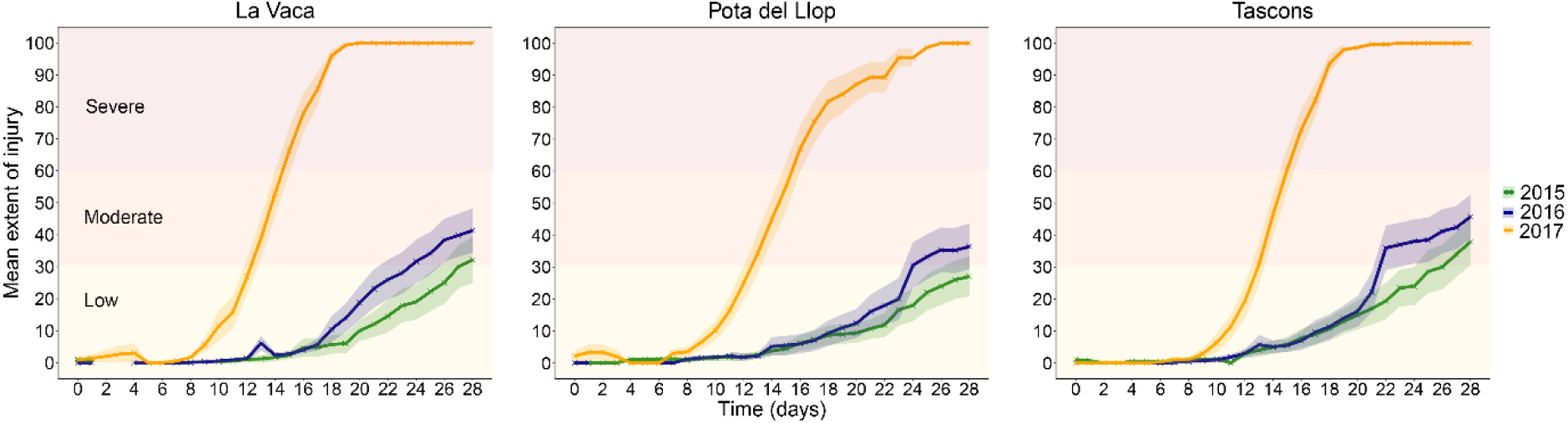
Average tissue necrosis (mean extent of injury ± SE) of *P. clavata* colonies during the 28 days of exposure in common garden experiments for La Vaca, Pota del Llop, and Tascons in 2015 (green), 2016 (blue) and 2017 (yellow). Mortality severity is highlighted in light yellow (low), orange (moderate), and red (severe).

### Individual fitness: modelling the response to thermal stress

The linear model 2 including the random factor ‘individual’ was retained (lowest statistically significant AIC = 710.88; df = 7 [Chi-square = 15.5, *p* < 0.001]). The ‘individual’ random factor had a significant effect, suggesting that each individual has a different baseline of resistance to necrosis. Regarding the fixed factors of the model, the deviance test showed that only the ‘year’ factor was significant (Table S2a), while *post-hoc* Tuckey test showed that significant differences were due to the year 2017 (Table S2b). The factor ‘year’ was contributing to 95.01% of the variance of the data, followed by the random factor ‘individual’ with a contribution of around 11 times less (4.1%). Finally, ‘population’ and ‘population-by-year interaction’ were non-significant and showed the lowest contributions: 0.5% and 0.39%, respectively.

### Genetic component: Genetic structure and individual heterosis

Results on heterozygosity, Hardy-Weinberg equilibrium, allelic richness, and linkage disequilibrium can be found in the Supplementary information (Tables S3 and S4, Figure S3). Three distinct genetic clusters matching the three populations in Medes islands were retrieved from the DAPC analysis with high mean membership probabilities over 84% for each cluster (Figure 3).

**Figure 3.**
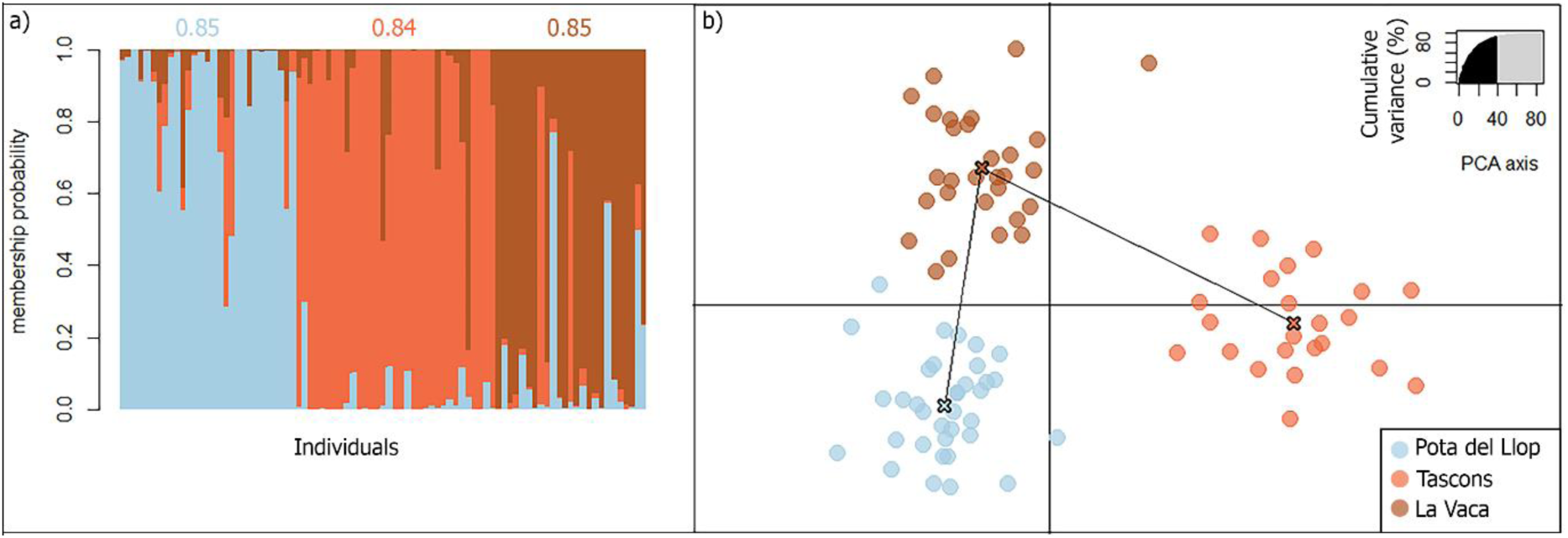
**a)** Individual membership probabilities are represented by a vertical line, where the different color segments indicate the individual proportion to each cluster (K=3) as estimated by the discriminant analysis of principal components (DAPC): Pota del Llop (light-blue), Tascons (orange) and La Vaca (brown). Mean membership probabilities are given above each colored cluster **b)** Scatter plot of the DAPC: each dot corresponds to one individual (N=87) from each of the three populations.

Overall, genetic differentiation was low but significant (global F_ST_=0.015, p<0.001). All pairwise F_ST_s were significant: La Vaca vs Pota del Llop (0.013; p<0.001), La Vaca vs Tascons (0.020; p<0.001), Pota del Llop vs Tascons (0.035; p<0.001). The analysis of site-specific F_ST_s suggested that Tascons was the most differentiated population (0.06, 95%HPDI: [0.030; 0.073]), followed by La Vaca (0.04, 95%HPDI: [0.022; 0.059]) and Pota del Llop (0.04, 95%HPDI: [0.021; 0.051]), albeit the differences were not significant (overlapping 95%HPDI; Table S5).

The standard multilocus heterozygosity (sMLH) ranged from 0.294 to 0.882 (Figure S4). The correlation between sMLH and “*nec-int*”, the proxy for individual fitness (random intercepts extracted from linear model 2), was not significant (r^2^ = 0.001, *p-value* = 0.66, Figure S5 and Table S6).

### Environmental component: thermal environments

Recent thermal history patterns, considered as June, July and August, revealed similar mean ± SD temperatures for the three years: 21.3°C ± 1.6 in 2015, 21.6°C ± 1.0 for 2016 and 21.8°C ± 1.3 for 2017 (Table S7). Concomitantly, extreme heat days (T23) were detected in all years during the summer season (Table S7). For 2015 and 2016, at 15m depth, maximum summer temperatures reached 24.8 and 23.5°C, respectively, and a low number of total extreme heat days exposure to ≥23°C was recorded (N=12 for 2015 and N=2 for 2016; Table S7, Figure S6a-b). The years 2015 and 2016 revealed several periods of anomalous high temperatures during the summer season, but no MHWs (Figure 4, Table S8). Interestingly, the year 2017 reported the highest maximum temperatures of 24.9 °C with a total of 19 days of exposure at extreme temperatures (≥23 °C), surpassing the thermal limit of *P. clavata* (Table S7, Figure S6). In addition, unlike for 2015 and 2016 where no MHW occurred, two MHWs occurred in 2017 with strong, and severe classifications from June to July and several heat spikes (Figure 4). The mean maximum temperature for these MHW was 24.1°C (Table S8).

**Figure 4.**
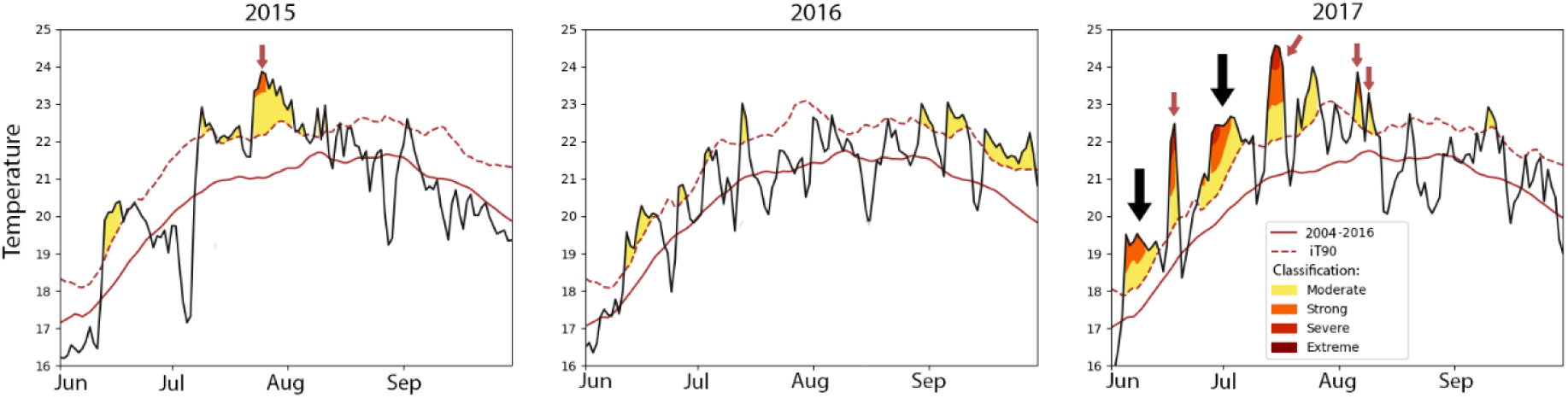
Daily mean temperature values recorded at 15m from June to September (Summer season) at Medes Islands with respect of the inter-annual climatological mean (red solid line) and 90^th^ percentile (iT90, red dotted line). Days below the red solid line were considered as “cool days”, whereas days above were considered as “warm days”. Following Hobday et al. 2018, marine heat waves and heat spikes (MHW and MHS) severity classification is as follows: “Moderate” (yellow), “Strong” (orange), “Severe” (red), and “Extreme” (dark-red). MHW are highlighted with a black arrow and MHS are highlighted with red arrows. Data taken from the T-MEDNet initiative (https://t-mednet.org/).

### Plastic component: ‘genotype-by-environment’ interactions and environmental sensitivity

Regarding the sensitivity analyses, environmental values (yearly mean PCA scores) were −0.08 ± 1.46 for 2015, 0.34 ± 1.55 for 2016, and 3.56 ± 0.55 for 2017. All individuals but one show positive slopes ranging between 0.16 and 1.65. Environmental sensitivity varied by a factor of 10 among individuals (Figure 5). Following our classification, genotypes were shared among hyper-sensitive and sensitive categories with no resistant genotypes (Figure 5).

**Figure 5.**
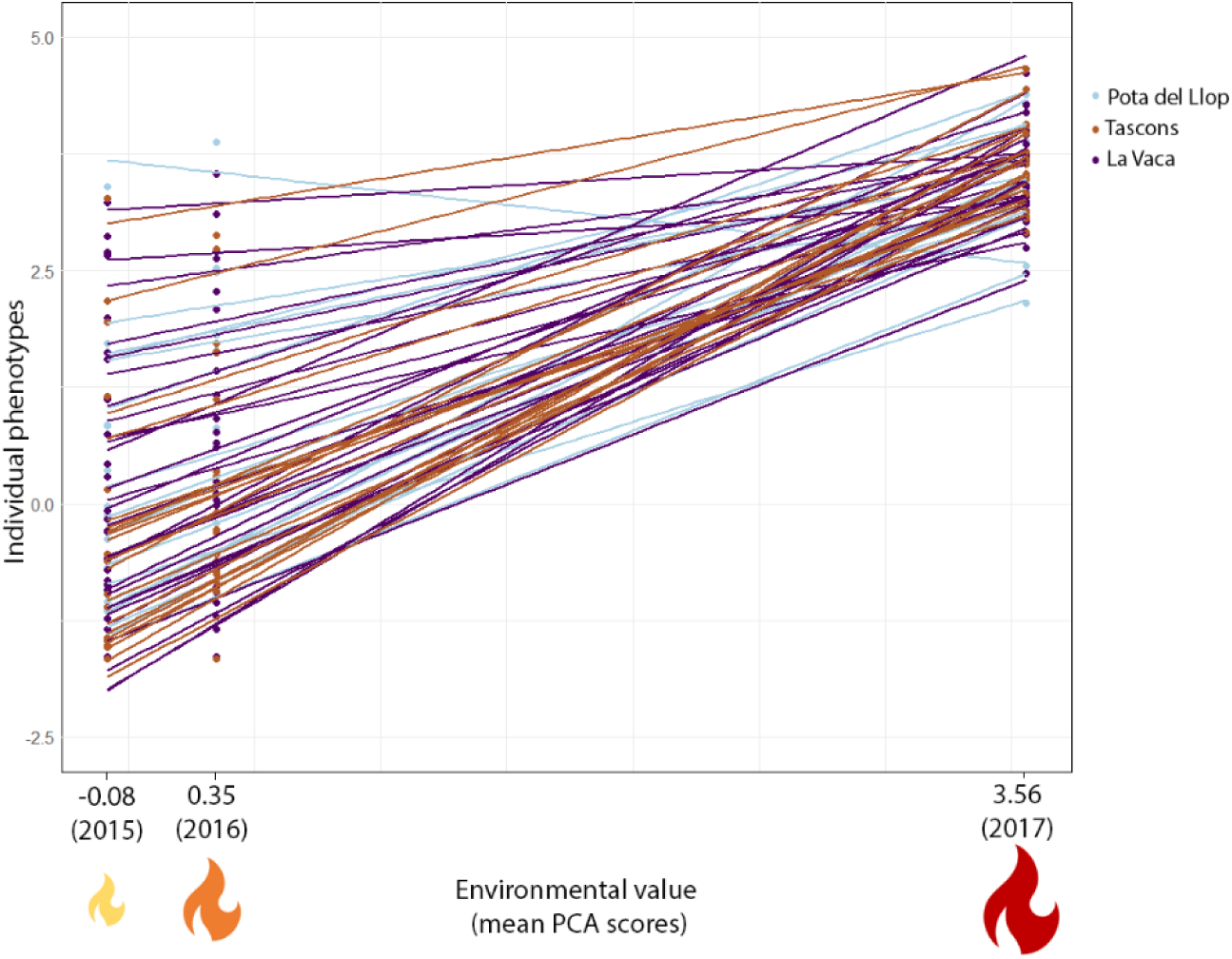
Sensitivity analyses and g*enotype-by-environment* interactions of *P. clavata* during 2015, 2016 and 2017 experiments. Individual ecological response (PCA scores) was plotted against each environment value (mean PCA score by year). The slope of the regression for each individual is considered as an estimator of the environmental sensitivity of the individual. Populations are displayed in colors: Pota del Llop (light-blue), Tascons (brown), and La Vaca (purple).

### Field survey of necrosis rates following 2018 and 2022 MMEs

We encountered 17, 22, and 21 colonies out of the 30 colonies initially marked in Pota del Llop, Tascons, and La Vaca, respectively. Six, three, and five colonies were not impacted (0% of necrosis) in La Vaca, Pota del Llop, and Tascons (Figure S6a). All the remaining colonies showed recent necrosis (>70%), albeit with different levels of impact (low, moderate, and severe, Figure S6a). Statistically significant differences were found in average tissue necrosis by population (F=4.295, p<0.018). Pota del Llop displayed the largest percentages of tissue necrosis (30.88% ± 37.22), followed by Tascons (28.63% ± 32.48) and La Vaca (7.38 % ± 6.25; Figure S6b). Finally, low impacts (<30%) were observed in all populations with seven, eight, and fifteen colonies in Pota del Llop, Tascons, and La Vaca (Figure S6a). This latter population did not report additional higher impacts while severe impacts (>60% tissue necrosis, including dead colonies) were observed in Pota del Llop and Tascons with four (three dead) and three (two dead) severely impacted colonies, respectively (Figure S6a).

## Discussion

We combined replicated experiments, population genetics, and an *in-situ* field survey to reveal the prominent influence of the environmental component (likely yearly variations in summer thermal regime) on the heterogeneity in ecological responses to thermal stress in *P. clavata*. The low influence of genetic and plastic components combined to the high environmental sensitivity of the tested genotypes point toward a dramatically low adaptive potential to recurrent MHWs. This “multi-event” perspective strengthens the recent call to carefully consider predictions on biodiversity evolution based on single-event experiments.

### The heterogeneity in the ecological response to thermal stress is mostly driven by the thermal regime during summer

The three populations of *Paramuricea clavata* showed high levels of tissue necrosis (moderate to severe mortality) at the end of each experiment confirming the species sensitivity to thermal stress (Crisci et al., 2017; Gómez-Gras et al., 2022). The ecological responses were similar between populations in 2015 and 2016 (moderate mortality) compared to 2017, in which colonies died before the end of the experiment (severe mortality). More than 95% of the variance in the ecological response was explained by the factor year (environmental component), while factors individual, population (genetic component), and the population-by-year interaction (plastic component) explained only 4.1%, 0.5%, and 0.39%, respectively. Factors individual and year (environmental component) were significant, which suggests different baselines of resistance to thermal stress among individuals and confirms the major environmental effect, mostly driven by the year 2017. This environmental effect was refined by the sensitivity and thermal regime analyses. First, the environmental value (mean phenotype) for 2017 was two to three times higher than the values for 2015 and 2016. Then, we reported positive regression slopes for almost all genotypes supporting an increased negative impact of environmental conditions (environmental sensitivity) from 2015 to 2017. Summer conditions during 2017 showed the largest number of MHS and MHWs compared to 2015 and 2016. Consequently, we posit that colonies of *P. clavata* were driven close to their physiological limits by the 2017 extreme summer conditions, which may have hampered any adjustment to thermal stress, whether genetic or plastic, during the experiment.

The relative impact of environmental, genetic, and plastic components in differential responses to ECEs has been screened in different species. For example, ubiquitous *population-by-environment* interactions (plastic component) have been detected in 172 species of plants (Matesanz & Ramírez-Valiente, 2019), but are lacking in others (Shao et al., 2022). Single event experiments with tropical corals identified local adaptation (*e.g.* Thomas et al., 2022) and adaptive plasticity (*e.g.* Drury et al., 2022) as drivers of differential bleaching responses. Here, we found relatively similar and high levels of necrosis among populations with a prevailing impact of the environmental component. These findings contrast to our previous studies based on “single event” experiments at larger spatial scales where differential ecological responses (distinct necrosis levels) were observed among populations in some cases (Arizmendi-Mejía, et al., 2015a; Arizmendi-Mejía et al., 2015b; Bonacolta et al., 2024; Crisci et al., 2017; Gómez-Gras et al., 2022). Two non-exclusive hypotheses relying on the spatial and temporal features of the experiments might explain this apparent discrepancy.

### Intraspecific differences in the response to thermal stress: does the geographic scale matter or are recent summer conditions overwhelming *P. clavata* physiological capacities?

The experiments conducted to date in *P. clavata* have considered a wide range of geographic distances, from local to regional (Arizmendi-Mejía et al., 2015b; Crisci et al., 2017) and inter-regional (Bonacolta et al., 2024; Gómez-Gras et al., 2022) scales. These experiments demonstrated population heterogeneity in ecological responses to thermal stress triggered by different drivers (*e.g.* genetic isolation, microorganisms). Considering that the impact of the genetic component on ecological responses can vary over spatial distances (*e.g.* Galloway & Fenster, 2000; Joshi et al., 2001), we hypothesize that the lack of significant population effect observed here can be related to the fine spatial scale of the experiment, which flattened the differences between populations. Contrary to previous experiments, we targeted three populations from similar habitats at the same depth range in a close spatial proximity (hundreds of meters). The potential for contrasted genetic make-up at such fine spatial scale is low (but see Ledoux et al., 2015; Richardson et al., 2014) as supported by the comparable levels of genetic isolation of the three populations (overlapping population-specific F_ST_s). Re-analyzing the different experiments in *P. Clavata* accounting for ecological or spatial distance between populations should allow to go further in this hypothesis.

Considering a temporal perspective, the discrepancy among experiments in *P. clavata* can result from an intensification of extreme climatic events from summer 2009 (Crisci et al., 2017) to summer 2019 (Gómez-Gras et al., 2022) and summer 2022 (this study; Rovira et al. 2024), which could have driven the colonies closer to their physiological limits in the later and warmer years. That is, the summer thermal regime previous to the first experiment (summer 2009, Crisci et al., 2017) was less stressful than the 2017 summer thermal regime (this study) allowing some colonies to face the 2009 experimental stress while colonies were totally swept by the 2017 experimental stress. Three main points support this hypothesis. First, the environmental sensitivity analysis considering 2015, 2016, and 2017 experiments clearly show a decrease in the variation of phenotypic responses among the individuals and an increase of the yearly environmental value between 2015/2016 and 2017. Individuals that showed relatively low necrosis in the first two years were as strongly impacted in 2017 (positive regression slopes) as individuals showing high necrosis during the first two experiments (regression slopes ~ 0). Second, necrosis was observed in a vast majority of the marked colonies in the three populations (>70%) during the field survey following the 2018 and 2022 MHWs. Third, one of the strongest mortality events ever observed was reported in 2022 (Estaque et al., 2023, Rovira et al. 2024) corroborating the rise in MHWs in the Mediterranean. Strengthening this hypothesis, the intensification of disastrous ECEs in the last decades is not peculiar to the Mediterranean (see Stillman, 2019). The frequency of bleaching events in tropical corals increased worldwide since 1980 (Hughes et al., 2018), with detrimental cumulative effects of heatwaves in the last ten years (Hughes et al., 2021).

### What’s next for shallow populations of *P. clavata*?

Our results question the persistence of the studied shallow populations of *P. clavata*. Both the experiments and field surveys conducted here supported a limited potential adaptability whether based on genetic or plastic component. Yet, we revealed some variability in the baseline ecological responses at the individual levels (4.1%). While this individual variability may have been seen as a hopeful “raw material” for adaptation to MHWs, the sensitivity analyses and the field survey showed how it was almost totally squeezed in 2017 (experiment) and 2022 (in the field). Dedicated studies combining population genomics, environmental, and mortality data are needed to further look for genome-environment associations, potential outlier loci involved in the differential responses and to estimate of the genomic offset of *P. clavata*. However, the lack of clear adaptive potential revealed here combined to the species life history traits (*e.g.* generation time >12 years, Coma et al., 1995) are such in a contrast with the current MHWs temporal dynamics that any evolutionary response seems compromised. The potential for adaptation is also questioned in tropical corals in which candidate genetic loci identified to date only show relatively elusive influence on heat stress tolerance (Fuller et al., 2020, but see Matz et al., 2020). In the same collapsing line, the absence of *population-by-environment* interactions suggest limited potential for evolutionary changes in adaptive plasticity (Sirovy et al., 2021). Recent studies point towards ecological memory, an increase in stress tolerance following previous exposure, as a key mechanism for coral acclimation to MHWs (Hackerott et al., 2021; Hughes et al., 2018). Yet, results are contrasted among species with a decrease in bleaching sensitivity following repeated heat stress in some species but not in others (Brown et al., 2023; Hughes et al., 2021). Our study allows first insights into *P. clavata* environmental memory. First, colony fragments used in a particular year were submitted to summer thermal conditions of the previous years. Yet, the worst ecological responses to thermal stress were observed during the last experiment in 2017 with high environmental sensitivity (positive slopes and lack of resistant genotypes in the sensitivity analyses). Then, most marked colonies showed necrosis during the field surveys following 2018 and 2022 MHWs events. These results question any increase in thermotolerance as expected with the ecological memory hypotheses and strengthen the limited potential for adaptation in *P. Clavata*.

## Conclusion

As temperature and frequency of ECEs continue to rise (Hughes et al., 2021; Garrabou et al., 2022), the evolution of biodiversity is more than ever a central concern for society. Adopting a “multi-event” perspective that combined replicated common-garden experiments in aquaria and mortality surveys in the field performed on the same colonies, our study points toward an inevitable collapse of many of the shallow populations of *P. clavata*. This collapse would emerge from a low to non-existent adaptive response, whether driven by genetic or plasticity, combined to a high environmental sensitivity and a potential intensification of MHWs. Considering the small spatial scale of our study, extrapolation at larger scale should be made cautiously. Yet, population collapses of *P. clavata* linked to recurrent MHWs have been observed in other Mediterranean regions (Garrabou et al., 2021; Gómez-Gras et al., 2021b). Moreover, field surveys following the 2022 MHW event in this and in other regions reported terrific mortality rates. Other populations from the same region showed a similar proportion of total affected colonies (70%) with almost 40% of the tissue necrosis (Rovira et al. 2024). Hundreds of km apart, populations until 20m depth displayed on average >80% of affected colonies and an increase by 142% of the degree of impact following the 2022 MHW compared to the previous MME in 2003 (Estaque et al., 2023). Worrying, this trend in *P. clavata* could likely be transposable to many of the Mediterranean habitat-forming and sessile species impacted by MHWs (Garrabou et al., 2022; Gómez-Gras et al., 2021a). We predict a shift in these species’ upper distribution limits, which will lead to a simplification of associated benthic communities hampering potentially related ecosystem functions and services (Gómez-Gras et al., 2021a).

This study echoes two recent calls regarding the future of marine diversity in the context of extreme climatic events. First, the impacts of ECEs on biodiversity should be studied from a temporal perspective, which accounts for their recurrence (Hughes et al., 2021). In this line, we suggest that predictions of biodiversity evolution based on “single event” approaches should be considered cautiously as they can be overly optimistic. Then, while conservation and restoration actions should be able to slow, and/or to some extent reverse, locally the collapsing trend followed by many marine habitat-forming species, immediate action on greenhouse gas emissions remains the only way to protect these species globally.

## Author’s contributions

**Sandra Ramirez-Calero:** Conceptualization, data curation, data analysis, writing original draft. **Daniel Gomez-Gras:** experimental setup, data collection, write – review and editing. **Aldo Barreiro**: data analysis and validation, write – review and editing. **Nathaniel Bensoussan**: data collection and validation, review and editing. **Laura Figuerola-Ferrando:** data collection, experimental setup, review and editing. **Marc Jou:** data visualization, review and editing. **A Lopez-Sanz:** data collection, experimental setup, review and editing, **Paula Lopez-Sendino:** data collection, experimental setup, review and editing. **Alba Medrano:** data collection, experimental setup, review and editing. **Ignasi Montero-Serra:** data collection, experimental setup, data analysis, review and editing. **Marta Pages-Escola:** data collection, experimental setup, review and editing. **Cristina Linares:** data collection, review and editing, funding acquisition. **Jean-Baptiste Ledoux:** conceptualization, data collection, experimental setup, data analysis, write – review and editing, funding acquisition. **Joaquim Garrabou:** conceptualization, data collection, experimental setup, data analysis, write – review and editing, funding acquisition.

## Acknowledgments

This work was financially supported by the European Union’s Horizon 2020 research and innovation programme [grants 689518—MERCES and SEP-210597628—FutureMARES], by MCIU/AEI/FEDER [RTI2018-095346-B-I00; HEATMED]. This research has also been funded by the project CORFUN (TED2021-131622B-I00) and by the Short-Term Scientific Mission (STSM) grant provided by the COST Action CA20102-MAF World (E-COST-GRANT-CA20102-f04dde3c). SRC was supported by the scholarship Doctorados en el exterior No. 906 de 2021 of Ministerio de Ciencia, Tecnología e Innovación of Colombia. JBL was supported by the strategic funding UIDB/04423/2020, UIDP/04423/2020 and 2021.00855.CEECIND through national funds provided by FCT-Fundaço para a Ciência e a Tecnologia. JG acknowledges the funding of the Spanish government through the ‘Severo Ochoa Centre of Excellence’ accreditation (CEX2019-000928-S). The authors would like to thank the technician staff at the Institute of Marine Sciences, Barcelona, for their support with the experimental setup and to the members of MedRecover research group (2021 SGR 01073) from the “Generalitat de Catalunya” for supporting fieldwork campaigns and laboratory procedures.

## Data availability statement

The data that support the findings of this study are openly available in the GitHub repository: “Pclavata_rec_MHW” at https://github.com/sandrarcr/Pclavata_rec_MHW.git (2024) including ecological data, linear models and genetic structure analysis. Temperature data and associated analysis can be found in “tmednetGUI” GitHub repository at https://github.com/Damyck/tMednet.git (2024), using the scripts: *surface_temperature.py*, *data_manager.py*, *user_interaction.py* and *excel_writer.py*. Finally, microsatellite loci information can be found under accession numbers GU386255–GU386265 in the Molecular Ecology Resources Database and GeneBank (https://10.1111/j.1755-0998.2010.02871.x).

## References

Ahrens, C. W., Challis, A., Byrne, M., Leigh, A., Nicotra, A. B., Tissue, D., & Rymer, P. (2021). Repeated extreme heatwaves result in higher leaf thermal tolerances and greater safety margins. New Phytologist, 232(3), 1212–1225. 10.1111/NPH.17640

Altwegg, R., Visser, V., Bailey, L. D., & Erni, B. (2017). Learning from single extreme events. Philosophical Transactions of the Royal Society B: Biological Sciences, 372(1723). 10.1098/RSTB.2016.0141

Arafeh-Dalmau, N., Schoeman, D. S., Montaño-Moctezuma, G., Micheli, F., Rogers-Bennett, L., Olguin-Jacobson, C., & Possingham, H. P. (2020). Marine heat waves threaten kelp forests. Science, 367(6478), 635. 10.1126/SCIENCE.ABA5244/ASSET/9E2E38F9-5365-4030-BC4F-707E0992A60F/ASSETS/SCIENCE.ABA5244.FP.PNG

Arizmendi-Mejía, R., Ledoux, J. B., Civit, S., Antunes, A., Thanopoulou, Z., Garrabou, J., & Linares, C. (2015). Demographic responses to warming: reproductive maturity and sex influence vulnerability in an octocoral. Coral Reefs, 34(4), 1207–1216. 10.1007/s00338-015-1332-9

Arizmendi-Mejía, R., Linares, C., Garrabou, J., Antunes, A., Ballesteros, E., Cebrian, E., Díaz, D., & Ledoux, J.-B. (2015). Combining Genetic and Demographic Data for the Conservation of a Mediterranean Marine Habitat-Forming Species. PLOS ONE, 10(3), e0119585. 10.1371/journal.pone.0119585

Bailey, L. D., & Van De Pol, M. (2016). Tackling extremes: challenges for ecological and evolutionary research on extreme climatic events. Journal of Animal Ecology, 85(1), 85–96. 10.1111/1365-2656.12451

Bay, R. A., & Palumbi, S. R. (2014). Multilocus adaptation associated with heat resistance in reef-building corals. Current Biology, 24(24), 2952–2956. 10.1016/j.cub.2014.10.044

Otero, M., Numa, C., Bo, M., Orejas, C., Garrabou, J., Cerrano, C., Kružic, P., Antoniadou, C., Aguilar, R., Kipson, S., Linares, C., Terrón-Sigler, A., Brossard, J., Kersting, D., Casado-Amezúa, P., García, S., Goffredo, S., Ocaña, O., Caroselli, E., … Özalp, B. (2017). Overview of the conservation status of Mediterranean anthozoa. Overview of the Conservation Status of Mediterranean Anthozoa. 10.2305/IUCN.CH.2017.RA.2.EN

Boavida, J., Assis, J., Silva, I., & Serrão, E. A. (2016). Overlooked habitat of a vulnerable gorgonian revealed in the Mediterranean and Eastern Atlantic by ecological niche modelling. Scientific Reports 6:1, 6(1), 1–9. 10.1038/srep36460

Bonacolta, A. M., Miravall, J., Gómez-Gras, D., Ledoux, J. B., López-Sendino, P., Garrabou, J., Massana, R., & del Campo, J. (2024). Differential apicomplexan presence predicts thermal stress mortality in the Mediterranean coral *Paramuricea clavata*. Environmental Microbiology, 26(1), e16548. 10.1111/1462-2920.16548

Brown, K. T., & Barott, K. L. (2022). The Costs and Benefits of Environmental Memory for Reef-Building Corals Coping with Recurring Marine Heatwaves. Integrative and Comparative Biology, 62(6), 1748–1755. 10.1093/ICB/ICAC074

Brown, K. T., Lenz, E. A., Glass, B. H., Kruse, E., McClintock, R., Drury, C., Nelson, C. E., Putnam, H. M., & Barott, K. L. (2023). Divergent bleaching and recovery trajectories in reef-building corals following a decade of successive marine heatwaves. Proceedings of the National Academy of Sciences of the United States of America, 120(52), e2312104120. 10.1073/PNAS.2312104120/SUPPL_FILE/PNAS.2312104120.SAPP.PDF

Cebrian, E., Linares, C., Marschal, C., & Garrabou, J. (2012). Exploring the effects of invasive algae on the persistence of gorgonian populations. Biological Invasions, 14(12), 2647–2656. 10.1007/S10530-012-0261-6/FIGURES/6

Chevin, L. M., Lande, R., & Mace, G. M. (2010). Adaptation, Plasticity, and Extinction in a Changing Environment: Towards a Predictive Theory. PLOS Biology, 8(4), e1000357. 10.1371/JOURNAL.PBIO.1000357

Coma, R., Ribes, M., Zabala, M., & Gili, J. M. (1995). Reproduction and cycle of gonadal development in the Mediterranean gorgonian *Paramuricea clavata*. Marine Ecology Progress Series, 117(1–3), 173–184. 10.3354/meps117173

Coma, R., Ribes, M., Zabala, M., & Gili, J. M. (1998). Growth in a Modular Colonial Marine Invertebrate. Estuarine, Coastal and Shelf Science, 47(4), 459–470. 10.1006/ECSS.1998.0375

Cramer, W., Guiot, J., Fader, M., Garrabou, J., Gattuso, J. P., Iglesias, A., Lange, M. A., Lionello, P., Llasat, M. C., Paz, S., Peñuelas, J., Snoussi, M., Toreti, A., Tsimplis, M. N., & Xoplaki, E. (2018). Climate change and interconnected risks to sustainable development in the Mediterranean. Nature Climate Change 8:11, 8(11), 972–980. 10.1038/s41558-018-0299-2

Crisci, C., Ledoux, J. B., Mokhtar-Jamaï, K., Bally, M., Bensoussan, N., Aurelle, D., Cebrian, E., Coma, R., Féral, J. P., La Rivière, M., Linares, C., López-Sendino, P., Marschal, C., Ribes, M., Teixidó, N., Zuberer, F., & Garrabou, J. (2017). Regional and local environmental conditions do not shape the response to warming of a marine habitat-forming species. Scientific Reports, 7(1), 1–13. 10.1038/s41598-017-05220-4

David, P. (1997). Modeling the genetic basis of heterosis: Tests of alternative hypotheses. Evolution, 51(4), 1049–1057. 10.1111/j.1558-5646.1997.tb03952.x

Dixon, G. B., Davies, S. W., Galina, A. V., Meyer, Eli., Bay, L. K., & Matz1, M. V. (2015). Genomic determinants of coral heat tolerance across latitudes. Science, 348(6242), 1460–1462. 10.1126/science.aaa7471

Drury, C., Dilworth, J., Majerová, E., Caruso, C., & Greer, J. B. (2022). Expression plasticity regulates intraspecific variation in the acclimatization potential of a reef-building coral. Nature Communications 2022 13:1, 13(1), 1–9. 10.1038/s41467-022-32452-4

Estaque, T., Richaume, J., Bianchimani, O., Schull, Q., Mérigot, B., Bensoussan, N., Bonhomme, P., Vouriot, P., Sartoretto, S., Monfort, T., Basthard-Bogain, S., Fargetton, M., Gatti, G., Barth, L., Cheminée, A., & Garrabou, J. (2023). Marine heatwaves on the rise: One of the strongest ever observed mass mortality event in temperate gorgonians. Global Change Biology, 29(22), 6159–6162. 10.1111/GCB.16931

Falconer, D. S., & Mackay, T. F. C. (1996). Introduction to quantitative genetics. Essex. UK: Longman Group. 448. Ed 4. Longmans Green, Harlow, Essex, UK.

Fitzpatrick, S. W., & Reid, B. N. (2019). Does gene flow aggravate or alleviate maladaptation to environmental stress in small populations? Evolutionary Applications, 12(7), 1402– 1416. 10.1111/EVA.12768

Foll, M., & Gaggiotti, O. (2006). Identifying the environmental factors that determine the genetic structure of populations. Genetics, 174(2), 875–891. 10.1534/genetics.106.059451

Fox, J., & Weisberg, S. (2019). R Companion to Applied Regression, Third edition. Sage, Thousand Oaks CA. Thousand Oaks CA: Sage., September 2012.

Frölicher, T. L., Fischer, E. M., & Gruber, N. (2018). Marine heatwaves under global warming. Nature 560:7718, 560(7718), 360–364. 10.1038/s41586-018-0383-9

Fuller, Z. L., Mocellin, V. J. L., Morris, L. A., Cantin, N., Shepherd, J., Sarre, L., Peng, J., Liao, Y., Pickrell, J., Andolfatto, P., Matz, M., Bay, L. K., & Przeworski, M. (2020). Population genetics of the coroal *Acropora millepora*: Toward genomic prediction of bleaching. Science, 369(6501). 10.1126/science.aba4674

Galloway, L. F., & Fenster, C. B. (2000). Population differentiation in an annual legume: local adaptation. Evolution, 54(4), 1173–1181. 10.1111/J.0014-3820.2000.TB00552.X

Garrabou, J., Gómez-Gras, D., Medrano, A., Cerrano, C., Ponti, M., Schlegel, R., Bensoussan, N., Turicchia, E., Sini, M., Gerovasileiou, V., Teixido, N., Mirasole, A., Tamburello, L., Cebrian, E., Rilov, G., Ledoux, J. B., Souissi, J. Ben, Khamassi, F., Ghanem, R., … Harmelin, J. G. (2022). Marine heatwaves drive recurrent mass mortalities in the Mediterranean Sea. Global Change Biology, 28(19), 5708–5725. 10.1111/gcb.16301

Garrabou, J., Ledoux, J.-B., Bensoussan, N., Gómez-Gras, D., & Linares, C. (2021). Sliding Toward the collapse of Mediterranean Coastal Marine Rocky Ecosystems. In Ecosystem Collapse and Climate Change (pp. 291–324). Springer, Cham. 10.1007/978-3-030-71330-0_11

Garrabou, J., Gómez-Gras, D., Ledoux, J.B., Linares, C., Bensoussan, N., López-Sendino, P., Bazairi, H., Espinosa, F., Ramdani, M., Grimes, S., Benabdi, M., Ben Souissi, J., Soufi, E., Khamassi, F., Ghanem, R., Ocaña, O., Ramos-Esplà, A., Izquierdo, A., Anton, A., … Haermelin, J. (2019). Collaborative database to track mass mortality events in the Mediterranean Sea. Front. Mar. Sci, 6, 707. 10.3389/fmars.2019.00707

Garrabou, J., Perez, T., Sartoretto, S., & Harmelin, J. (2001). Mass mortality event in red coral Corallium rubrum populations in the Provence region (France, NW Mediterranean). Marine Ecology Progress Series, 217, 263–272. 10.3354/meps217263

Genin, A., Levy, L., Sharon, G., Raitsos, D. E., & Diamant, A. (2020). Rapid onsets of warming events trigger mass mortality of coral reef fish. Proceedings of the National Academy of Sciences of the United States of America, 117(41), 25378–25385. 10.1073/PNAS.2009748117/-/DCSUPPLEMENTAL

Gómez-Gras, D., Bensoussan, N., Ledoux, J. B., López-Sendino, P., Cerrano, C., Ferretti, E., Kipson, S., Bakran-Petricioli, T., Serrao, E. A., Paulo, D., Coelho, M. A. G., Pearson, G. A., Boavida, J., Montero-Serra, I., Pagès-Escolà, M., Medrano, A., López-Sanz, A., Milanese, M., Linares, C., & Garrabou, J. (2022). Exploring the response of a key Mediterranean gorgonian to heat stress across biological and spatial scales. Scientific Reports, 12(1), 21064. 10.1038/s41598-022-25565-9

Gómez-Gras, D., Linares, C., Dornelas, M., Madin, J. S., Brambilla, V., Ledoux, J. B., López-Sendino, P., Bensoussan, N., & Garrabou, J. (2021). Climate change transforms the functional identity of Mediterranean coralligenous assemblages. Ecology Letters, 24(5), 1038–1051. 10.1111/ele.13718

Gómez-Gras, D., Linares, C., López-Sanz, A., Amate, R., Ledoux, J. B., Bensoussan, N., Drap, P., Bianchimani, O., Marschal, C., Torrents, O., Zuberer, F., Cebrian, E., Teixidó, N., Zabala, M., Kipson, S., Kersting, D. K., Montero-Serra, I., Pagès-Escolà, M., Medrano, A., Frleta-Valić, M., Frleta-Valić, M., Frleta-Valić, M., Garrabou, J. (2021). Population collapse of habitat-forming species in the Mediterranean: a long-term study of gorgonian populations affected by recurrent marine heatwaves. Proceedings of the Royal Society B: Biological Sciences, 288(1965), 20212384. 10.1098/rspb.2021.2384

Hackerott, S., Martell, H. A., & Eirin-Lopez, J. M. (2021). Coral environmental memory: causes, mechanisms, and consequences for future reefs. Trends in Ecology and Evolution, 36(11), 1011–1023. 10.1016/j.tree.2021.06.014

Hobday, A.J., E.C.J. Oliver, A. Sen Gupta, J.A. Benthuysen, M.T. Burrows, M.G. Donat, N.J. Holbrook, P.J. Moore, M.S. Thomsen, T. Wernberg, and D.A. Smale. 2018. Categorizing and naming marine heatwaves. Oceanography 31(2):162–173, 10.5670/oceanog.2018.205

Hobday, A. J., Alexander, L. V., Perkins, S. E., Smale, D. A., Straub, S. C., Oliver, E. C. J., Benthuysen, J. A., Burrows, M. T., Donat, M. G., Feng, M., Holbrook, N. J., Moore, P. J., Scannell, H. A., Sen Gupta, A., & Wernberg, T. (2016). A hierarchical approach to defining marine heatwaves. Progress in Oceanography, 141, 227–238. 10.1016/j.pocean.2015.12.014

Hothorn, T., Bretz, F., & Westfall, P. (2008). Simultaneous inference in general parametric models. In Biometrical Journal,50(3). 10.1002/bimj.200810425

Hughes, T. P., Anderson, K. D., Connolly, S. R., Heron, S. F., Kerry, J. T., Lough, J. M., Baird, A. H., Baum, J. K., Berumen, M. L., Bridge, T. C., Claar, D. C., Eakin, C. M., Gilmour, J. P., Graham, N. A. J., Harrison, H., Hobbs, J. P. A., Hoey, A. S., Hoogenboom, M., Lowe, R. J., … Wilson, S. K. (2018). Spatial and temporal patterns of mass bleaching of corals in the Anthropocene. Science, 359(6371), 80–83. 10.1126/science.aan8048

Hughes, T. P., Baird, A. H., Bellwood, D. R., Card, M., Connolly, S. R., Folke, C., Grosberg, R., Hoegh-Guldberg, O., Jackson, J. B. C., Kleypas, J., Lough, J. M., Marshall, P., Nyström, M., Palumbi, S. R., Pandolfi, J. M., Rosen, B., & Roughgarden, J. (2003). Climate change, human impacts, and the resilience of coral reefs. Science, 301(5635), 929–933. 10.1126/SCIENCE.1085046/ASSET/7D57E257-3B37-4EC8-8514-9F0CA04132F9/ASSETS/GRAPHIC/SE3231777004.JPEG

Hughes, T. P., Kerry, J. T., Connolly, S. R., Álvarez-Romero, J. G., Eakin, C. M., Heron, S. F., Gonzalez, M. A., & Moneghetti, J. (2021). Emergent properties in the responses of tropical corals to recurrent climate extremes. Current Biology, 31(23), 5393–5399.e3. 10.1016/j.cub.2021.10.046

Hughes, T. P., Kerry, J. T., Connolly, S. R., Baird, A. H., Eakin, C. M., Heron, S. F., Hoey, A. S., Hoogenboom, M. O., Jacobson, M., Liu, G., Pratchett, M. S., Skirving, W., & Torda, G. (2018). Ecological memory modifies the cumulative impact of recurrent climate extremes. Nature Climate Change 9:1, 9(1), 40–43. 10.1038/s41558-018-0351-2

Husson, B., Lind, S., Fossheim, M., Kato-Solvang, H., Skern-Mauritzen, M., Pécuchet, L., Ingvaldsen, R. B., Dolgov, A. V., & Primicerio, R. (2022). Successive extreme climatic events lead to immediate, large-scale, and diverse responses from fish in the Arctic. Global Change Biology, 28(11), 3728–3744. 10.1111/GCB.16153

Jacox, M. G., Alexander, M. A., Bograd, S. J., & Scott, J. D. (2020). Thermal displacement by marine heatwaves. Nature 584:7819, 584(7819), 82–86. 10.1038/s41586-020-2534-z

Jombart, T., Devillard, S., & Balloux, F. (2010). Discriminant analysis of principal components: a new method for the analysis of genetically structured populations. BMC Genetics, 11. 10.1186/1471-2156-11-94

Joshi, J., Schmid, B., Caldeira, M. C., Dimitrakopoulos, P. G., Good, J., Harris, R., Hector, A., Huss-Danell, K., Jumpponen, A., Minns, A., Mulder, C. P. H., Pereira, J. S., Prinz, A., Scherer-Lorenzen, M., Siamantziouras, A. S. D., Terry, A. C., Troumbis, A. Y., & Lawton, J. H. (2001). Local adaptation enhances performance of common plant species. Ecology Letters, 4(6), 536–544. 10.1046/J.1461-0248.2001.00262.X

Lê, S., Josse, J., & Husson, F. (2008). FactoMineR: An R package for multivariate analysis. Journal of Statistical Software, 25(1). 10.18637/jss.v025.i01

Ledoux, J. B., Aurelle, D., Bensoussan, N., Marschal, C., Féral, J. P., & Garrabou, J. (2015). Potential for adaptive evolution at species range margins: Contrasting interactions between red coral populations and their environment in a changing ocean. Ecology and Evolution, 5(6), 1178–1192. 10.1002/ece3.1324

Ledoux, J. B., Frleta-Valić, M., Kipson, S., Antunes, A., Cebrian, E., Linares, C., Sánchez, P., Leblois, R., & Garrabou, J. (2018). Postglacial range expansion shaped the spatial genetic structure in a marine habitat-forming species: Implications for conservation plans in the Eastern Adriatic Sea. Journal of Biogeography, 45(12), 2645–2657. 10.1111/jbi.13461

Leung, J. Y. S., Connell, S. D., & Russell, B. D. (2017). Heatwaves diminish the survival of a subtidal gastropod through reduction in energy budget and depletion of energy reserves. Scientific Reports 2017 7:1, 7(1), 1–8. 10.1038/s41598-017-16341-1

Linares, C., Coma, R., Mariani, S., Díaz, D., Hereu, B., & Zabala, M. (2008). Early life history of the Mediterranean gorgonian *Paramuricea clavata*: Implications for population dynamics. Invertebrate Biology, 127(1), 1–11. 10.1111/j.1744-7410.2007.00109.x

Logan, C. A., Dunne, J. P., Eakin, C. M., & Donner, S. D. (2014). Incorporating adaptive responses into future projections of coral bleaching. Global Change Biology, 20(1), 125–139. 10.1111/GCB.12390

Malyshev, A. V., Arfin Khan, M. A. S., Beierkuhnlein, C., Steinbauer, M. J., Henry, H. A. L., Jentsch, A., Dengler, J., Willner, E., & Kreyling, J. (2016). Plant responses to climatic extremes: within-species variation equals among-species variation. Global Change Biology, 22(1), 449–464. 10.1111/gcb.13114

Matesanz, S., & Ramírez-Valiente, J. A. (2019). A review and meta-analysis of intraspecific differences in phenotypic plasticity: Implications to forecast plant responses to climate change. In Global Ecology and Biogeography, 28(11), 1682–1694. 10.1111/geb.12972

Matz, M. V., Treml, E. A., & Haller, B. C. (2020). Estimating the potential for coral adaptation to global warming across the Indo-West Pacific. Global Change Biology, 26(6), 3473–3481. 10.1111/GCB.15060

Maxwell, S. L., Butt, N., Maron, M., McAlpine, C. A., Chapman, S., Ullmann, A., Segan, D. B., & Watson, J. E. M. (2019). Conservation implications of ecological responses to extreme weather and climate events. In Diversity and Distributions, 25(4), 613–625. 10.1111/ddi.12878

Merilä, J., & Hendry, A. P. (2014). Climate change, adaptation, and phenotypic plasticity: The problem and the evidence. Evolutionary Applications, 7(1), 1–14. 10.1111/EVA.12137

Mokhtar-Jamaï, K., Pascual, M., Ledoux, J. B., Coma, R., Féral, J. P., Garrabou, J., & Aurelle, D. (2011). From global to local genetic structuring in the red gorgonian *Paramuricea clavata*: The interplay between oceanographic conditions and limited larval dispersal. Molecular Ecology, 20(16), 3291–3305. 10.1111/j.1365-294X.2011.05176.x

Montero-Serra, I., Garrabou, J., Doak, D. F., Ledoux, J. B., & Linares, C. (2019). Marine protected areas enhance structural complexity but do not buffer the consequences of ocean warming for an overexploited precious coral. Journal of Applied Ecology, 56(5), 1063–1074. 10.1111/1365-2664.13321

Pansch, C., Scotti, M., Barboza, F. R., Al-Janabi, B., Brakel, J., Briski, E., Bucholz, B., Franz, M., Ito, M., Paiva, F., Saha, M., Sawall, Y., Weinberger, F., & Wahl, M. (2018). Heat waves and their significance for a temperate benthic community: A near-natural experimental approach. Global Change Biology, 24(9), 4357–4367. 10.1111/GCB.14282

Parmesan C, Root TL, & Willig MR. (2000). Impacts of extreme weather and climate on terrestrial biota. Bull Am Meteor Soc, 81(3), 443–450. 10.1175/1520-0477(2000)081 <0443:IOEWAC>2.3.CO;2

Pinsky, M. L., Eikeset, A. M., McCauley, D. J., Payne, J. L., & Sunday, J. M. (2019). Greater vulnerability to warming of marine versus terrestrial ectotherms. Nature, 569(7754), 108–111. 10.1038/S41586-019-1132-4

Ponti, M., Perlini, R. A., Ventra, V., Grech, D., Abbiati, M., & Cerrano, C. (2014). Ecological shifts in Mediterranean coralligenous assemblages related to gorgonian forest loss. PloS One, 9(7). 10.1371/JOURNAL.PONE.0102782

Ponti, M., Turicchia, E., Ferro, F., Cerrano, C., & Abbiati, M. (2018). The understorey of gorgonian forests in mesophotic temperate reefs. Aquatic Conservation: Marine and Freshwater Ecosystems, 28(5), 1153–1166. 10.1002/AQC.2928

R Core Team. (2022). R: A language and environment for statistical computing. R Foundation for Statistical Computing, Vienna, Austria. https://www.R-project.org/

Raymond, M., & Rousset, F. (1995). GENEPOP (version 1.2): Population genetics software for exact tests and ecumenicism. Journal of Heredity, 86, 248–249. 10.1093/oxfordjournals.jhered.a111573

Regan, C. E., & Sheldon, B. C. (2023). Phenotypic plasticity increases exposure to extreme climatic events that reduce individual fitness. Global Change Biology. 10.1111/gcb.16663

Reusch, T. B. H. (2014). Climate change in the oceans: evolutionary versus phenotypically plastic responses of marine animals and plants. Evolutionary Applications, 7(1), 104–122. 10.1111/EVA.12109

Richardson, J. L., Urban, M. C., Bolnick, D. I., & Skelly, D. K. (2014). Microgeographic adaptation and the spatial scale of evolution. Trends in Ecology & Evolution, 29(3), 165–176. 10.1016/J.TREE.2014.01.002

Risso, A., Risso, A., Plée, V., & Prêtre, J. G. (1826). Histoire naturelle des principales productions de l’Europe méridionale et particulièrement de celles des environs de Nice et des Alpes Maritimes. In *Histoire naturelle des principales productions de l’Europe méridionale et particulièrement de celles des environs de Nice et des Alpes Maritimes*. Chez F.-G. Levrault, libraire. 10.5962/bhl.title.58984

Rousset, F. (2008). GENEPOP’007: A complete re-implementation of the GENEPOP software for Windows and Linux. Molecular Ecology Resources, 8(1), 103–106. 10.1111/j.1471-8286.2007.01931.x

Rovira, G., Capdevila, P., Zentner, Y., Margarit, N., Ortega, J., Casals, D., Figuerola-Ferrando, L., Aspillaga, E., Medrano, A., Pagès-Escolà, M., Hereu, B., Garrabou, J., Linares, C. (2024). When resilience is not enough: 2022 extreme marine heatwave threatens climatic refugia for a habitat-forming Mediterranean octocoral. J. Animal Ecology. *In press*.

Scheiner, S. M. (1993). Genetics and Evolution of Phenotypic Plasticity. Annu. Rev. Ecol Syst, 24, 35–68. 10.1146/ANNUREV.ES.24.110193.000343

Shao, J., Li, G., Li, Y., & Zhou, X. (2022). Intraspecific responses of plant productivity and crop yield to experimental warming: A global synthesis. Science of The Total Environment, 840, 156685. 10.1016/J.SCITOTENV.2022.156685

Sirovy, K. A., Johnson, K. M., Casas, S. M., La Peyre, J. F., & Kelly, M. W. (2021). Lack of genotype-by-environment interaction suggests limited potential for evolutionary changes in plasticity in the eastern oyster, *Crassostrea virginica*. Molecular Ecology, 30(22), 5721–5734. 10.1111/mec.16156

Smale, D. A., & Wernberg, T. (2013). Extreme climatic event drives range contraction of a habitat-forming species. Proceedings of the Royal Society B: Biological Sciences, 280(1754). 10.1098/RSPB.2012.2829

Smale, D. A., Wernberg, T., Oliver, E. C. J., Thomsen, M., Harvey, B. P., Straub, S. C., Burrows, M. T., Alexander, L. V., Benthuysen, J. A., Donat, M. G., Feng, M., Hobday, A. J., Holbrook, N. J., Perkins-Kirkpatrick, S. E., Scannell, H. A., Sen Gupta, A., Payne, B. L., & Moore, P. J. (2019). Marine heatwaves threaten global biodiversity and the provision of ecosystem services. Nature Climate Change 2019 9:4, 9(4), 306–312. 10.1038/s41558-019-0412-1

Smith, K. E., Burrows, M. T., Hobday, A. J., King, N. G., Moore, P. J., Sen Gupta, A., Thomsen, M. S., Wernberg, T., & Smale, D. A. (2023). Biological Impacts of Marine Heatwaves. Annual Review of Marine Science, 15, 119–145. 10.1146/annurev-marine-032122-121437

Stillman, J. H. (2019). Heat waves, the new normal: Summertime temperature extremes will impact animals, ecosystems, and human communities. Physiology, 34(2), 86–100. 10.1152/PHYSIOL.00040.2018/ASSET/IMAGES/LARGE/PHY0021904600006.JPEG

Stoffel, M. A., Esser, M., Kardos, M., Humble, E., Nichols, H., David, P., & Hoffman, J. I. (2016). inbreedR: an R package for the analysis of inbreeding based on genetic markers. Methods in Ecology and Evolution, 7(11), 1331–1339. 10.1111/2041-210X.12588

Thomas, L., Underwood, J. N., Rose, N. H., Fuller, Z. L., Richards, Z. T., Dugal, L., Grimaldi, C. M., Cooke, I. R., Palumbi, S. R., & Gilmour, J. P. (2022). Spatially varying selection between habitats drives physiological shifts and local adaptation in a broadcast spawning coral on a remote atoll in Western Australia. Science Advances, 8(17), 9185. 10.1126/SCIADV.ABL9185/SUPPL_FILE/SCIADV.ABL9185_TABLES_S1_TO_S9.ZIP

Ummenhofer, C., & Meehl, G. (2017). Extreme weather and climate events with ecological relevance: a review. Philosophical Transactions of the Royal Society B: Biological Sciences, 372(20160135). 10.1098/rstb.2016.0135

van Bergen, E., Dallas, T., DiLeo, M. F., Kahilainen, A., Mattila, A. L. K., Luoto, M., & Saastamoinen, M. (2020). The effect of summer drought on the predictability of local extinctions in a butterfly metapopulation. Conservation Biology, 34(6), 1503–1511. 10.1111/COBI.13515

Venables, W. N., & Ripley, B. D. (2002). Modern Applied Statistics with S. Fourth edition by. Springer New York, NY.

von Schuckmann, K., Le Traon, P. Y., Smith, N., Pascual, A., Djavidnia, S., Gattuso, J. P., Grégoire, M., Nolan, G., Aaboe, S., Aguiar, E., Álvarez Fanjul, E., Alvera-Azcárate, A., Aouf, L., Barciela, R., Behrens, A., Belmonte Rivas, M., Ben Ismail, S., Bentamy, A., Borgini, M., … Zuo, H. (2019). Copernicus Marine Service Ocean State Report, Issue 3. Journal of Operational Oceanography, 12(sup1), S1–S123. 10.1080/1755876X.2019.1633075

Weir, B. S., & Cockerham, C. C. (1984). Estimating F-Statistics for the Analysis of Population Structure. Evolution, 38(6), 1358. 10.2307/2408641

Zentner, Y., Rovira, G., Margarit, N., Ortega, J., Casals, D., Medrano, A., Pagès-Escolà, M., Aspillaga, E., Capdevila, P., Figuerola-Ferrando, L., Riera, J. L., Hereu, B., Garrabou, J., & Linares, C. (2023). Marine protected areas in a changing ocean: Adaptive management can mitigate the synergistic effects of local and climate change impacts. Biological Conservation, 282, 110048. 10.1016/J.BIOCON.2023.110048

