## Supplementary information for "Recurrent extreme climatic events are driving gorgonian populations to local extinction: low adaptive potential to marine heatwaves"

In this Supplementary Information, we provide further details regarding:

- the “*Common garden set-up*” including Figure S1,
- the “*Phenotypic response: survival analysis, individual fitness and modeling the response to thermal stress*” including the methods to compute the PCAs used to define the proxy for the phenotypic response to thermal stress (and corresponding Figure S2)
- the *“DNA extraction and microsatellites characteristics”* including the protocols for DNA extraction, microsatellites amplification (Tables S1) and genotyping as well as methods used to computed rarefied allelic richness, heterozygosity and estimator of FIS.
- the “*Individual fitness: modelling the response to thermal stress* including Table S2 showing the Analysis of deviance and Turkey post hoc tests.
- the “*Genetic component: Genetic structure and individual heterosis*” including results of the microsatellites quality check and genetic diversity (Table S3), linkage desequilibrium (Figure S3) and Hardy-Weinberg equilibrium (Table S4). We also provide the results of the site-specific Fst (Table S5) and details regarding the analyses of the heterosis, which includes the standardized multilocus heterozygosity (sMLH) values (Figure S4) and the correlation between sMLH and the phenotypic responses (*nec-int*) (Figure S5 and Table S6).
- the “*Environmental component: summer thermal environments”* including a characterization of the temperature records at Medes Islands for yearly summer seasons at 15m (Table S7), the numbers of extreme heat days of exposure (Figure S8) and of Marine Heatwaves (MHW) (Table S8).
- The *“Field survey of necrosis rates following 2018-2022 MMEs”* including a description of the level of mortality in the three populations (Figure S6).

*Common garden set-up*

The experiment was conducted at the Aquarium Experimental Zone at the ICM-CSIC (Spain). Upon arrival from the field, colony fragments were mechanically fixed to experimental plates as described in Crisci et al. (2017) (Figure S1).

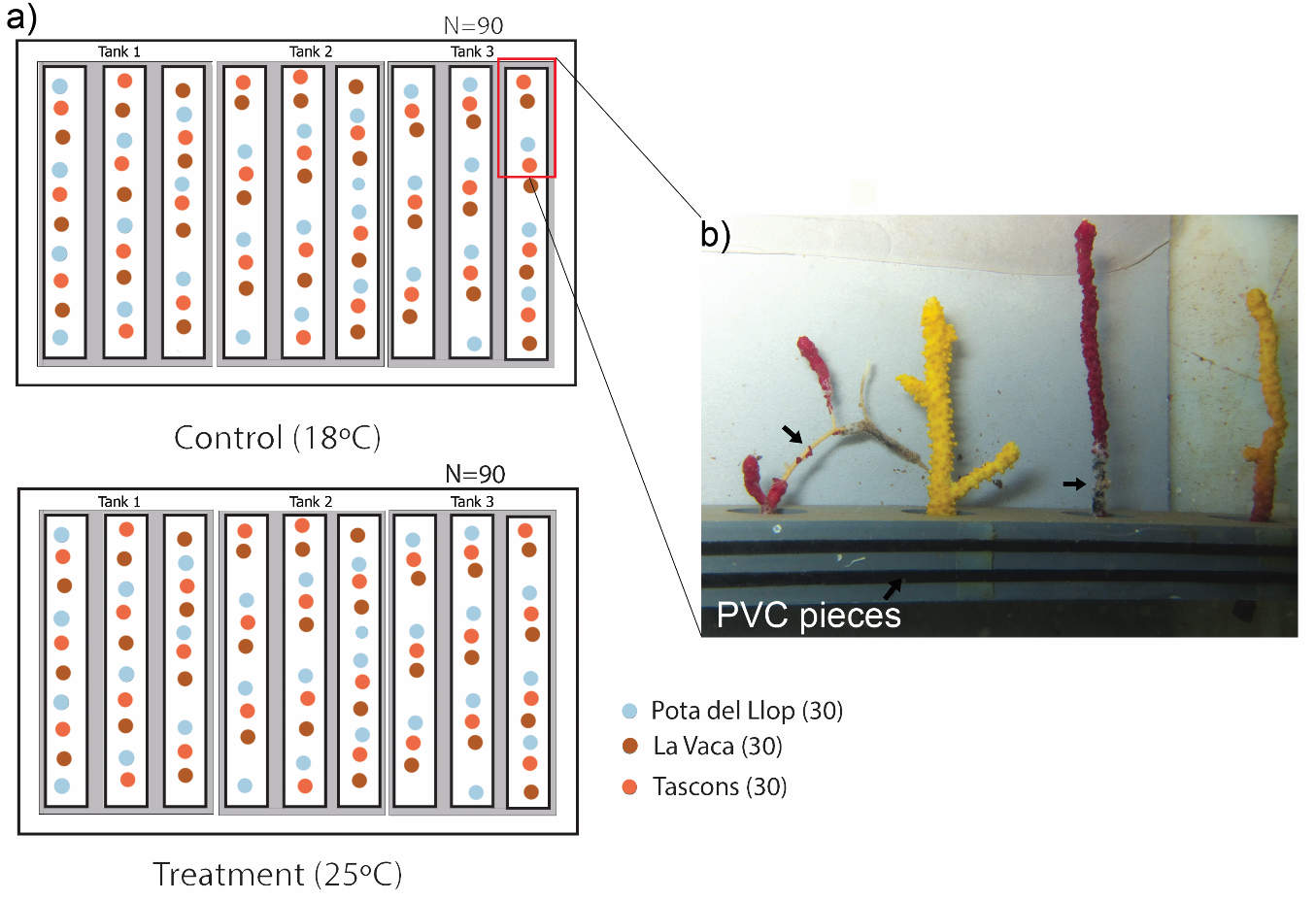

**Figure S1**. a) Experimental setup including tagged colonies after collection at Medes islands. Ninety fragments were randomly distributed in an open-water system aquarium setup. b) Individual fragments fixed to PVC rectangular pieces during experimental exposure. Black arrows highlight initial signs of tissue necrosis caused by high temperatures.

The common garden experiment involved two aquaria sets: Control and Treatment, each involving three replicate tanks (70 L each) with 10 different individuals from each population in each tank (30 individuals per population and experimental conditions). In addition, a buffer tank (also 70 L) was used to control seawater temperature in both Control and Treatment tanks. Each buffer tank was supplied with 50 μm sand-filtered Mediterranean seawater pumped from 15 m depth directly into the experimental tanks, functioning as an open system. All colonies were acclimated for one week at 17-18°C. In the Control set, seawater temperature was maintained at 16–18 °C during the whole experiment. In the Treatment set, the heat stress consisted of a stepwise temperature increase from 18 to 25°C over 3 days (25°C ± 0.2; Fig. S1). Upon reaching 25 °C, thermal conditions were kept constant for 28 days. To monitor the temperature in each experimental tank throughout the experiment, HOBO temperature data loggers were used (accuracy 0.2 C, resolution 0.02 C, temperature record every 5 min). One submersible water pump was used in each tank. Feeding was carried out three times per week during the 2nd and 6th days each week with 3 ml of a liquid mixture of particles between 10 and 450 μM in size (Bentos Nutrition Marine Active Supplement, Maim, Vic, Spain), and a tablet of frozen cyclops (Ocean nutrition, Antwerp, Belgium) on the 4th day (Crisci et al., 2017; Gómez-Gras et al., 2019).

*Phenotypic response: survival analysis, individual fitness and modeling the response to thermal stress*

To model the response to thermal stress across years and populations, we computed a Principal Component Analysis (PCA) with individuals and years as case and the percentage of necrosis per day as variables. A first exploratory PCA explained 50.29% of the variance and was strongly correlated with percentage of necrosis from day 10th to 28th (Figure S2a). To improve the fitness of the data, days 1 to 9 were removed from the data set. Thus, the second PCA explained the 75% of the variance for the remaining days (Figure S2b). All PCAs were performed with *PCA()* function from *FactoMineR* R package (Lê et al., 2008). Scores from the second PCA were considered as a proxy for individual fitness as they were correlated to the baseline percentages of necrosis.

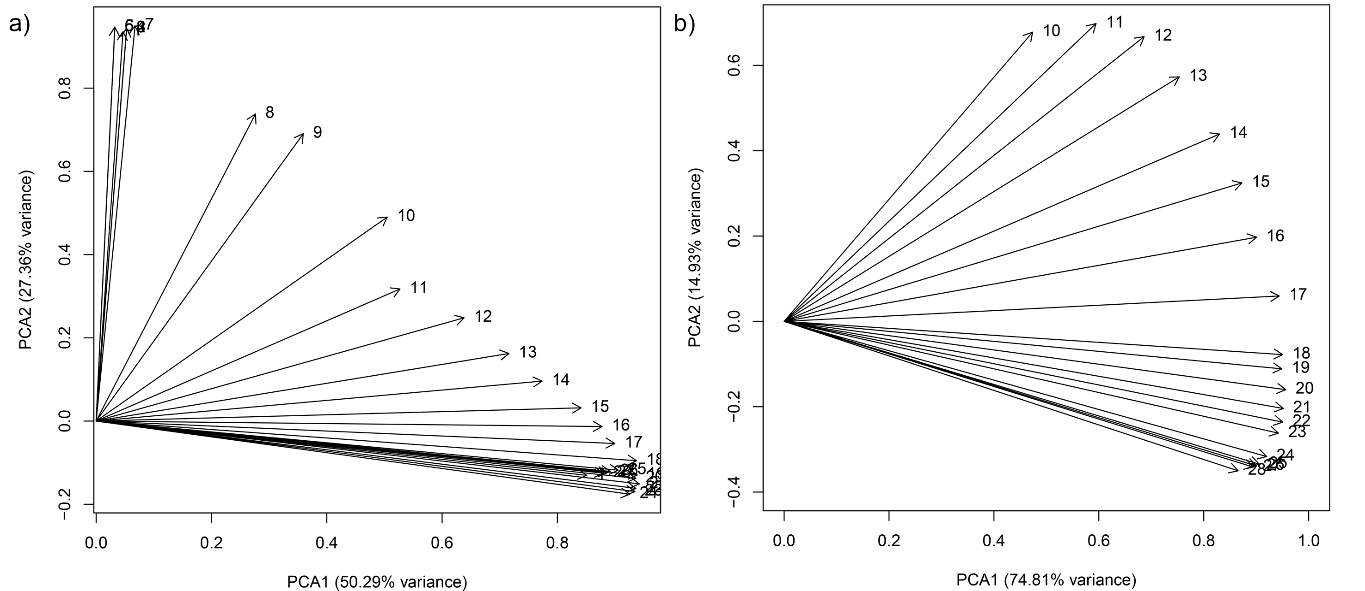

**Figure S2.** Principal Component Analysis (PCA) to obtain proxy for individual fitness. a) first exploratory PCA explaining 50% of the variance in the data (including all days), b) second posterior PCA explaining 75% of the variance in the data (10^th^ day to 28^th^).

*DNA extraction and microsatellites characteristics*

Total genomic DNA of 87 individuals was extracted following a salting out procedure adjusted from (Miller et al., 1988). These individuals were genotyped with 17 microsatellites previously isolated for the species. Seven microsatellites (Pcla 9, Pcla 10, Pcla 12, Pcla 14, Pcla 17, Pcla 81, Pcla a) were amplified as in Agell et al., 2009; Aurelle et al., 2010; Ledoux et al., 2018). The remaining 10 were developed for this study and amplified with PCR conditions as described in Table S1. PCR products were genotyped on an ABI3730 sequencer at the Plateforme Génome Transcriptome de Bordeaux (France; http://www4.bordeaux-aquitaine.inra.fr/pgtb) using the size standard ladder GeneScan 600-LIZ (Applied Biosystems). STRand v.2.4.59 was used for allele scoring (UC Davis Veterinary Genetics Laboratory). The genotyping procedures were validated by re-extracting and re-amplifying around ten percent of the samples.

**Table S1**. Primer sequences, PCR conditions and genetic characteristics of specific microsatellites for *Paramuricea clavata*.

| Locus name | Primer sequence (5’-3’) | Repeat motif | Number of cycles | T_a_(ºC) | Allele size range (bp) | Reference |
| --- | --- | --- | --- | --- | --- | --- |
| Pcla 20 | F: TCGCCCAAAGATGAGTAGGG  R: GTGTCTGCACATTTATGTTCGC | (AATA)_8_ | 35 | 56 | 245-269 | *Not available* |
| Pcla 21 | F: CGGCGATGTAATTCAGCCAC  R: AGTCCTACAAGTTTTCGATATTTCTTG | (TGCG)_8_ | 35 | 56 | 236-287 | *Not available* |
| Pcla 22 | F: TCAACTAAAGGTAATCTGAATGGC  R: ATCGCATGACACGTTCTTATC | (TAT)_8_ | 35 | 56 | 145-226 | *Not available* |
| Pcla 23 | F: GATATCTGGGCACCATCAGC  R: AGTTACGCAGATGTAAAGCTACC | (ATA)_8_ | 35 | 56 | 174-211 | *Not available* |
| Pcla 24 | F: ACTGCCTCGTAACATAACAACG  R: ATGGTCAAGAACCTTATGCAAG | (TTGA)_15_ | 35 | 56 | 179-233 | *Not available* |
| Pcla 25 | F: CAAGACAGTTCCTGCATGGTC  R: ACGCCTTATGTTTTCAGATACGAG | (TGC)_9_ | 35 | 56 | 197-223 | *Not available* |
| Pcla 26 | F: GACAAATCCGTAGGGACTAACAC  R: CCCCGGCACTTTGTAATAAGC | (TAA)_14_ | 35 | 56 | 182-197 | *Not available* |
| Pcla 27 | F: GCCCGTCTTTTATCCATGCC  R: CAACGTACGAGGATGTTGCC | (ATCA)_8_ATCC  (ATCA)_8_ | 35 | 56 | 163-239 | *Not available* |
| Pcla 28 | F: GTCATAGAGAACTATGCATGGCG  R: TTGGCTACTGATTCACAAACG | (TTAT)_7_ | 35 | 56 | 191-204 | *Not available* |
| Pcla 29 | F: TAAACACCATGCCAAGTCCG  R: GTTGCACGTATGATGTAACTGG | (ATGT)_12_ | 35 | 56 | 175-257 | *Not available* |

We used MICRO-CHECKER 2.2.3 (van Oosterhout et al., 2004) to test scoring errors due to stutters and large allele dropout. Several R packages were used to estimate missing data and genetic diversity statistics. First, microsatellites data genotyped for 90 individuals initially were stored into a *genind* object created with *adegenet* package (Goudet & Jombart, 2022; Jombart, 2008). The percentage of missing data was estimated using the function *info_table()* from *poppr* package (Kamvar et al., 2015) in order to remove potential defective loci, using a total cutoff of 1% for each locus within the whole population of Medes Islands. Rarefied allelic richness, gene diversity (Ht), observed (Ho) and expected heterozygosities (He) and Fis were assessed for each locus and population using the packages *hierfstat* (Goudet & Jombart, 2022) and *pegas* (Paradis, 2010). Hardy-Weinberg equilibrium (HWE) was assessed using *hw.test()* in *pegas* and *Genepop* (Raymond & Rousset, 1995; Rousset, 2008). Linkage disequilibrium (LD) was tested using *ia()* by calculating the association index of alleles among different loci in each populations implemented in *poppr* (Agapow & Burt, 2001; Brown et al., 1980).

*Individual fitness: modelling the response to thermal stress.*

The linear model 2 including the random factor ‘individual’ was retained (lowest statistically significant AIC = 710.88; df = 7 [Chi-square = 15.5, *p* < 0.001]). The ‘individual’ random factor had a significant effect, suggesting that each individual has a different baseline of resistance to necrosis. Regarding the fixed factors of the model, the deviance test showed that only the “year” factor was significant (Table S2a), while *post-hoc* Tuckey test showed that significant differences were due to the year 2017 (Table S2b).

**Table S2.** a) Analysis of deviance for intercepts using ‘population’ and ‘year’ as predictor variables while ‘individual’ as random. b) Tukey *post-hoc* for ‘Year’. Significant values are highlighted in bold.

| a) |  |  |  |
| --- | --- | --- | --- |
| Model factor | Chi-squared | *df* | *p*-value |
| Population | 0.89 | 2 | 0.64 |
| Year | 528.42 | 2 | **< 0.001** |

| b) |  |  |  |
| --- | --- | --- | --- |
| Comparison | Mean difference | *z* value | *p-value* |
| 2016 - 2015 | 0.35 | 1.9 | 0.13 |
| 2017 - 2015 | 3.64 | 20.3 | **< 0.001** |
| 2017 - 2016 | 3.29 | 19.2 | **< 0.001** |

*Genetic component: Genetic structure and individual heterosis.*

Using MICRO-CHECKER 2.2.3, we did not detected errors linked to stutters or large allele dropouts. Three loci with high proportions of missing data in the population were eliminated: Pcla20 (62%), Pcla10 (~21%) and Pcla29 (~12%). Rarefied allelic richness (Ar) ranged from 3 (Pcla23 in Vaca, Pota and Tascons) to 12.08 (Pcla14 in Pota) (Table S3). Observed heterozygosities (Ho) ranged from 0.44 (Pcla22) to 0.89 (Pcla9) (Table S3). Similar values of gene diversity (Ht) ranged from 0.51 (Pcla24) to 0.9 (Pcla14) (Table S3)

**Table S3.** Rarefied allelic richness (Ar) estimates for 14 microsatellite loci per population. Overall statistics for all loci including observed heterozygosity (Ho), gene diversities (Hs) and Fis.

|  | Allelic richness | | |  |  |  |
| --- | --- | --- | --- | --- | --- | --- |
|  | La Vaca | Pota del Llop | Tascons | Ho | Hs | Fis |
| Pcla21 | 5.19 | 5.15 | 5.37 | 0.72 | 0.72 | 0.00 |
| Pcla22 | 4.95 | 4.49 | 4.70 | 0.44 | 0.66 | 0.33 |
| Pcla.23 | 3.00 | 3.00 | 3.00 | 0.59 | 0.59 | 0.00 |
| Pcla.24 | 3.72 | 3.00 | 3.80 | 0.57 | 0.51 | -0.13 |
| Pcla.a | 6.79 | 4.82 | 5.33 | 0.79 | 0.73 | -0.08 |
| Pcla12 | 11.13 | 9.13 | 10.23 | 0.80 | 0.82 | 0.03 |
| Pcla14 | 11.63 | 12.08 | 11.06 | 0.86 | 0.90 | 0.05 |
| Pcla17 | 5.01 | 3.64 | 3.99 | 0.40 | 0.57 | 0.30 |
| Pcla25 | 4.89 | 3.32 | 3.81 | 0.49 | 0.52 | 0.07 |
| Pcla26 | 3.00 | 3.88 | 3.56 | 0.57 | 0.64 | 0.10 |
| Pcla27 | 5.89 | 6.62 | 6.59 | 0.64 | 0.64 | 0.00 |
| Pcla28 | 4.50 | 3.80 | 4.79 | 0.51 | 0.56 | 0.09 |
| Pcla81 | 5.89 | 5.64 | 4.81 | 0.67 | 0.70 | 0.05 |
| Pcla9 | 11.00 | 9.80 | 10.34 | 0.89 | 0.89 | 0.00 |

Index of association of alleles among different loci (r_d_) used to test the occurrence of linkage disequilibrium ranged from -0.2 – 0.4 in la Vaca, -0.1 – 0.1 in Tascons and from -0.2 - 0.4 in Pota del Llop (Figure S3). No linkage disequilibrium was consistent among the three populations. We therefore retained all the microsatellites

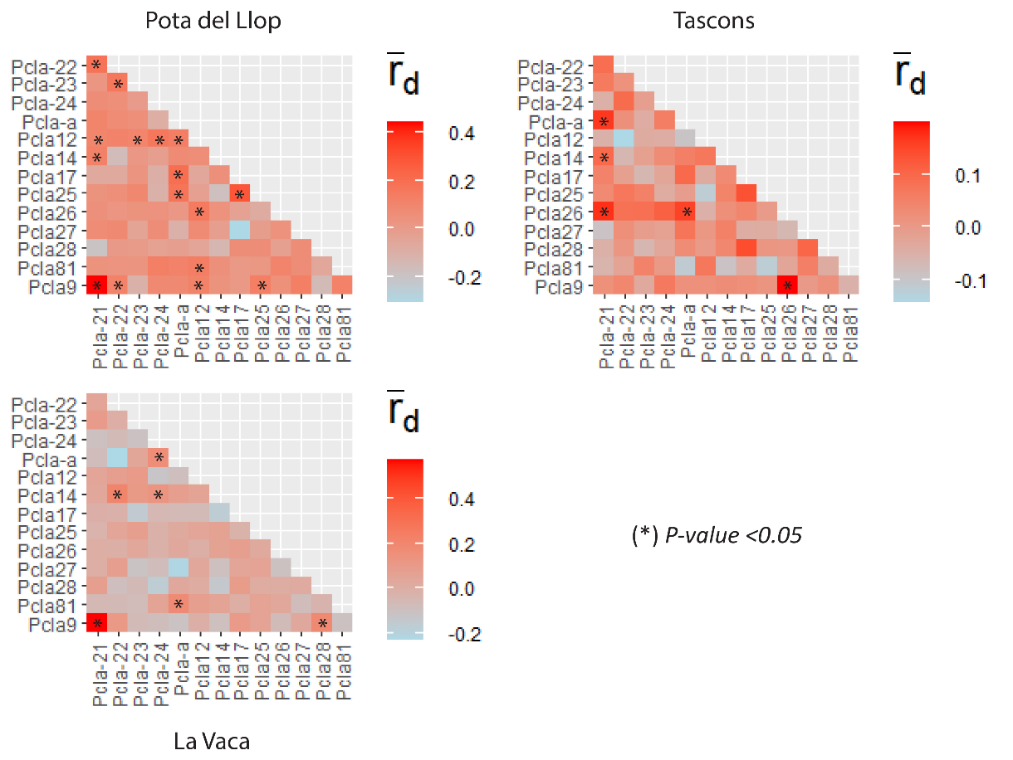

**Figure S3.** Pairwise linkage disequilibrium among loci in *P. clavata* populations. Significant deviations are highlighted with * (p<0.05).

Deviations from Hardy-Weinberg Equilibrium (HWE) were observed in different loci in each of the three populations (p<0.05, Table S4).

**Table S4**. Chi-squared tests of Hardy-Weinberg Equilibrium (HWE) for Pota del Llop, Tascons and La Vaca. Significant values are shown in bold.

|  | Pota del Llop | | | Tascons | | | La Vaca | | |
| --- | --- | --- | --- | --- | --- | --- | --- | --- | --- |
|  | *chi^2* | df | *p-value* | *chi^2* | df | *p-value* | *chi^2* | df | *p-value* |
| Pcla-21 | 9.11 | 15 | 0.872 | 13.41 | 15 | 5.71e-01 | 8.371 | 15 | 9.08e-01 |
| Pcla-22 | 21.84 | 10 | **0.016** | 3.31 | 15 | 9.99e-01 | 46.88 | 10 | **9.95e-07** |
| Pcla-23 | 1.08 | 3 | 0.782 | 2.41 | 3 | 4.93e-01 | 0.95 | 3 | 8.13e-01 |
| Pcla-24 | 3.17 | 6 | 0.788 | 1.62 | 3 | 6.56e-01 | 2.80 | 6 | 8.33e-01 |
| Pcla a | 24.72 | 36 | 0.922 | 14.21 | 15 | 5.10e-01 | 7.68 | 15 | 9.36e-01 |
| Pcla 12 | 124.55 | 120 | 0.370 | 77.30 | 78 | 5.01e-01 | 145.50 | 78 | **5.56e-06** |
| Pcla 14 | 93.99 | 105 | 0.771 | 175.89 | 136 | **1.20e-02** | 88.47 | 78 | 1.96e-01 |
| Pcla 17 | 25.52 | 15 | **0.043** | 11.60 | 6 | 7.15e-02 | 11.64 | 6 | 7.053e-02 |
| Pcla 25 | 37.12 | 15 | **0.001** | 10.54 | 6 | 1.04e-01 | 3.68 | 6 | 7.20e-01 |
| Pcla 26 | 5.60 | 3 | 0.132 | 46.66 | 6 | **2.19e-08** | 3.20 | 6 | 7.83e-01 |
| Pcla 27 | 43.68 | 21 | **0.003** | 21.44 | 21 | 4.32e-01 | 29.79 | 28 | 3.73e-01 |
| Pcla 28 | 8.87 | 10 | 0.545 | 3.92 | 6 | 6.88e-01 | 14.96 | 10 | 1.34e-01 |
| Pcla 81 | 13.61 | 21 | 0.886 | 11.42 | 15 | 7.22e-01 | 5.17 | 10 | 8.80e-01 |
| Pcla 9 | 56.68 | 91 | 0.998 | 57.33 | 66 | 7.68e-01 | 75.08 | 66 | 2.08e-01 |

The site-specific F_ST_ were computed in GESTE to estimate the relative impact of genetic drift on the differentiation of the considered site relative to the remaining ones. Results are shown in Table S5.

**Table S5**. Site-specific Fst and 95% HPDI in GESTE.

|  | Local Fst | 95% HPDI |
| --- | --- | --- |
| Pota del Llop | 0.036 | [0.022;0.052] |
| Tascons | 0.051 | [0.030;0.073] |
| Vaca | 0.040 | [0.023;0.059] |

We looked for heterosis in the response to thermal stress testing the correlation between the values of *nec-int* as proxy for individual fitness and the standardized individual multilocus heterozygosities (*sMLH)* computed using the R package *InbreedR()* (Stoffel et al., 2016). The slope of the linear model [*lm(nec-int ~ sMLH)*] was compared to its null distribution obtained with a Monte-Carlo permutation test with 10,000 permutations. The standard multilocus heterozygosity (sMLH) ranged from 0.294 to 0.882 (Figure S4). The correlation between sMLH and “*nec-int*”, the proxy for individual fitness (random intercepts extracted from linear model 2), was not significant (r^2^ = 0.001, *p-value* = 0.66, Figure S5 and Table S6).

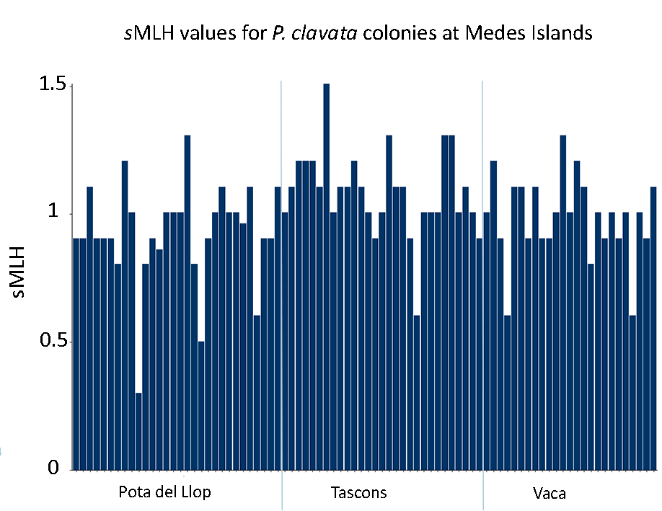

**Figure S4**. Multilocus heterozygosities (sMLH) were calculated for each *P. clavata* colony used during the experiment.

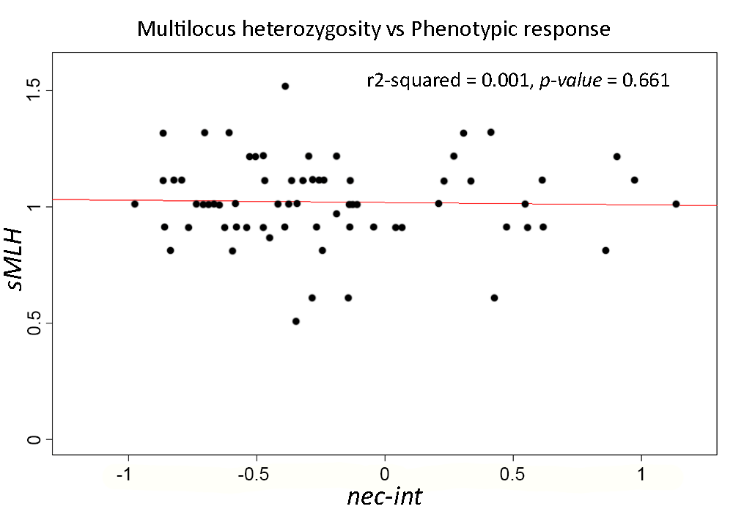

**Figure S5.** Linear regression model of *P. clavata* individual heterozygosities (MLH, genotypic response) function of individual *nec-int* (phenotypic responses). R^2^ = 0.001, *p>*0.66. Values of *nec-int* range between -1 to 1, representing phenotypic response of individuals developing less (resistant) or more (sensitive) necrosis.

**Table S6**. Parameters of the predictive *lm()* for *P. clavata*. Estimate, coefficient estimate of explanatory variables: *nec-int* and MLH; SE, test statistic (t value) and probability (*p*-value <0.05). Significant *p*-values are highlighted in bold.

| Variable | Estimate | SE | *t* value | | *p*-value |
| --- | --- | --- | --- | --- | --- |
| (Intercept) | 0.996 | 0.022 | 45.576 | **<2e-16** | |
| *nec-int* | -0.018 | 0.040 | -0.441 | 0.661 | |

*Environmental component: summer thermal environments*

Below, we characterize the recent thermal history patterns for the three years (2015, 2016, 2017).

**Table S7.** Temperature records at Medes Islands for yearly summer seasons at 15m. Number of records (N), Mean and Standard Deviation (Std), Maximum (Max) and Minimum (Min) temperatures are reported. The total number of days with sustained temperatures surpassing 23ºC and 24ºC are also shown (Ndays).

| **Year** | **N** | **Mean** | **Std** | **Max** | **Min** | **Ndays>=23** | **Ndays>=24** |
| --- | --- | --- | --- | --- | --- | --- | --- |
| 2017 | 3 | 21.28 | 1.96 | 24.90 | 15.03 | 14 | 5 |
| 2016 | 3 | 20.58 | 1.73 | 23.47 | 15.41 | 2 | 0 |
| 2015 | 3 | 20.71 | 2.24 | 24.75 | 15.13 | 11 | 1 |

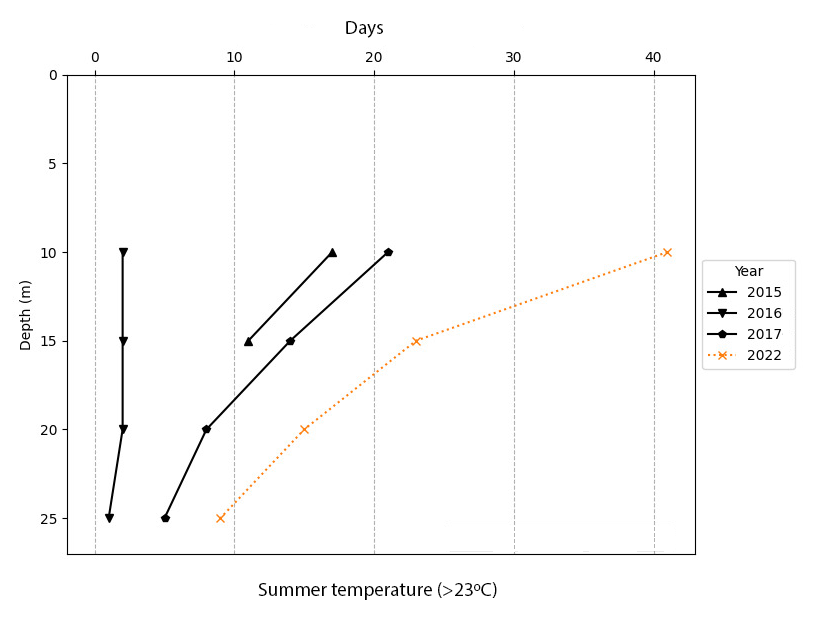

**Figure S6**. Number of extreme heat days of exposure for *P. clavata* during 2015-2017. a) for days at 23ºC and b) for days at 24ºC.

**Table S8**. Number of Marine Heatwaves (MHW) reported for the summer months of the periods of study. Values calculated from fixed thermometers at 15 at Medes islands.

| *Date (y/m/d)* | *Depth (m)* | *Duration (days)* | *Max intensity (ºc)* | *Cumulative intensity (ºc day)* | *Mean intensity (ºc)* | *Mean temperature* |
| --- | --- | --- | --- | --- | --- | --- |
| *2017-06-28* | 15 | 7 | 2.33 | 15.14 | 2.16 | 22.46 |
| *2017-07-13* | 15 | 5 | 3.26 | 13.85 | 2.77 | 24.07 |

*Field survey of necrosis rates following 2018-2022 MMEs:*

We surveyed by scuba diving the percentage of tissue necrosis *in situ* for the same individuals used in the experiments. This survey was done in October 2022 and the results are presented here and in the main text (Figure S6).

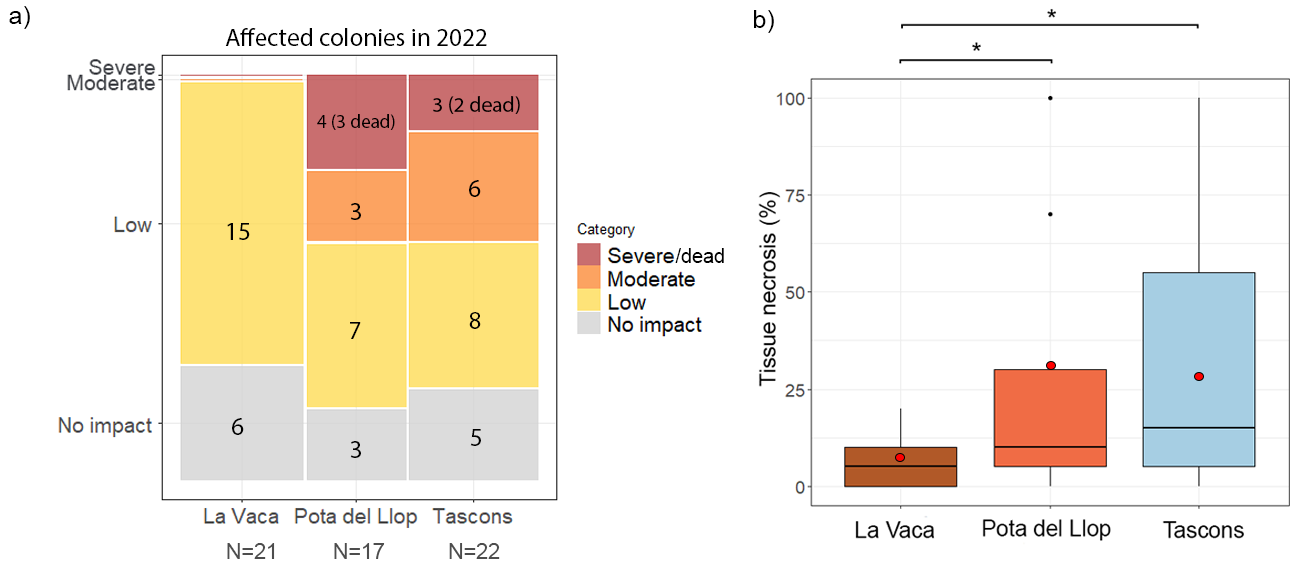

**Figure S6.** a) Mosaic plot representing impact severity according to tissue necrosis levels in colonies of *P. clavata* during 2022 for each population. Squares are proportional to the number of colonies that fell into each category: no impact (gray; <10% tissue necrosis), low (yellow; 10-30% tissue necrosis), moderate (orange; 30-60% tissue necrosis), and severe impacts (red; >60% tissue necrosis and including the number of dead colonies). b) Percentages of tissue necrosis for marked colonies of *P. clavata* during 2022. Boxes represent the interquartile range (IQR), encompassing the middle 50% of the data for La Vaca (brown), Pota del Llop (light-blue) and Tascons (orange). Black horizontal lines for each box represent the median, indicating the central tendency of the data, while red dots indicate the mean. Significant differences among populations are represented with (*) at the top of each comparison.

**References**

Agapow, P. M., & Burt, A. (2001). Indices of multilocus linkage disequilibrium. *Molecular Ecology Notes*, *1*(1–2), 101–102. <https://doi.org/10.1046/j.1471-8278.2000.00014.x>

Agell, G., Rius, M., & Pascual, M. (2009). Isolation and characterization of eight polymorphic microsatellite loci for the Mediterranean gorgonian Paramuricea clavata. In *Conservation Genetics,* 10(6) 2025–2027). <https://doi.org/10.1007/s10592-009-9885-1>

Aurelle, D., Baker, A. J., Bottin, L., Brouat, C., Caccone, A., Chaix, A., Dhakal, P., Ding, Y., Duplantier, J. M., Fiedler, W., Fietz, J., Fong, Y., Forcioli, D., Freitas, T. R. O., Gunnarsson, G. H., Haddrath, O., Hadziabdic, D., Hauksdottir, S., Havill, N. P., … Yuan, Q. Y. (2010). Permanent genetic resources added to the molecular ecology resources database 1 February 2010-31 March 2010. *Molecular Ecology Resources*, *10*(4), 751–754. <https://doi.org/10.1111/j.1755-0998.2010.02871.x>

Brown, A. H. D., Feldman, M. W., & Nevo, E. (1980). Multilocus Structure of Natural Populations of HORDEUM SPONTANEUM. *Genetics*, *96*(2), 523–536. <https://doi.org/10.1093/GENETICS/96.2.523>

Crisci, C., Ledoux, J. B., Mokhtar-Jamaï, K., Bally, M., Bensoussan, N., Aurelle, D., Cebrian, E., Coma, R., Féral, J. P., La Rivière, M., Linares, C., López-Sendino, P., Marschal, C., Ribes, M., Teixidó, N., Zuberer, F., & Garrabou, J. (2017). Regional and local environmental conditions do not shape the response to warming of a marine habitat-forming species. *Scientific Reports*, *7*(1), 1–13. <https://doi.org/10.1038/s41598-017-05220-4>

Gómez-Gras, D., Linares, C., de Caralt, S., Cebrian, E., Frleta-Valić, M., Montero-Serra, I., Pagès-Escolà, M., López-Sendino, P., & Garrabou, J. (2019). Response diversity in Mediterranean coralligenous assemblages facing climate change: Insights from a multispecific thermotolerance experiment. *Ecology and Evolution*, *9*(7), 4168–4180. <https://doi.org/10.1002/ece3.5045>

Goudet, J., & Jombart, T. (2022). hierfstat: Estimation and Tests of Hierarchical F-Statistics. R package version 0.5-11. In *R Core Team*.

Jombart, T. (2008). Adegenet: A R package for the multivariate analysis of genetic markers. *Bioinformatics*, *24*(11), 1403–1405. <https://doi.org/10.1093/bioinformatics/btn129>

Kamvar, Z. N., Brooks, J. C., & Grünwald, N. J. (2015). Novel R tools for analysis of genome-wide population genetic data with emphasis on clonality. *Frontiers in Genetics*, *6*(JUN), 208. <https://doi.org/10.3389/fgene.2015.00208>

Lê, S., Josse, J., & Husson, F. (2008). FactoMineR: An R package for multivariate analysis. *Journal of Statistical Software*, *25*(1). <https://doi.org/10.18637/jss.v025.i01>

Miller, S. A., Dykes, D. D., & Polesky, H. F. (1988). A simple salting out procedure for extracting DNA from human nucleated cells. *Nucleic Acids Research1*, *16*(3), 1215.

Paradis, E. (2010). Pegas: An R package for population genetics with an integrated-modular approach. *Bioinformatics*, 26(3) 419–420). <https://doi.org/10.1093/bioinformatics/btp696>

Raymond, M., & Rousset, F. (1995). GENEPOP (version 1.2): Population genetics software for exact tests and ecumenicism. *Journal of Heredity*, *86*, 248–249.

Rousset, F. (2008). GENEPOP’007: A complete re-implementation of the GENEPOP software for Windows and Linux. *Molecular Ecology Resources*, *8*(1), 103–106. <https://doi.org/10.1111/j.1471-8286.2007.01931.x>

Stoffel, M. A., Esser, M., Kardos, M., Humble, E., Nichols, H., David, P., & Hoffman, J. I. (2016). inbreedR: an R package for the analysis of inbreeding based on genetic markers. *Methods in Ecology and Evolution*, *7*(11), 1331–1339. <https://doi.org/10.1111/2041-210X.12588>

van Oosterhout, Cock Hutchinson, W., Wills, D., & Shipley, P. (2004). MICRO-CHECKER: software for identifying and correcting genotyping errors in microsatellite data. *Molecular Ecology Notes*, *4*, 535–538. <http://moscow.sci-hub.bz/f9cd0df5426111b33248b635261f44e6/>
